# Learning-specific remodeling of the hippocampal palmitoylome

**DOI:** 10.64898/2026.05.15.725211

**Authors:** Agata Pytyś, Athira Nataraj, Rafał Polowy, Rabia Ijaz, Alonso Cerdeño-Arévalo, Lucía Murillo-Hernández, Ángela Fontán-Lozano, Jakub Włodarczyk, Robert Kuba Filipkowski, Rebeca Mejías, Kasia Radwanska, Tomasz Wójtowicz

## Abstract

Protein S-palmitoylation is a reversible lipid modification that regulates protein trafficking, membrane association, and synaptic signaling, yet its role in learning-induced neuronal plasticity remains incompletely understood. Here, we investigated how spatial learning remodels the hippocampal palmitoylome in rats trained in the Morris water maze using short-term (STT; one session, 15 trials; probe at 1 h) or long-term (LTT; four sessions over four days; probe at 24 h) paradigms. Palmitoylated proteins were profiled by acyl-biotin exchange coupled with tandem mass tag labeling and LC-MS/MS. We identified 5,260 proteins, including 763 palmitoylated species. Spatial learning induced extensive remodeling of protein S-palmitoylation, with markedly greater changes after STT than LTT. Using yoked controls, we distinguished palmitoylation changes associated with learning the hidden platform location from those induced by general behavioral experience. Comparison of trained and yoked animals identified 186 and 62 differentially palmitoylated proteins after STT and LTT, respectively, whereas yoked animals also exhibited extensive changes relative to naïve controls, demonstrating that behavioral experience alone substantially reshapes the hippocampal palmitoylome. Functional enrichment revealed that STT preferentially engaged pathways related to synaptic transmission, cytoskeletal remodeling, GTPase signaling, and cellular metabolism, whereas LTT was associated with protein translation and synaptic organization. Site-specific analysis identified numerous previously unreported palmitoylation sites. Hierarchical analysis further identified diacylglycerol lipase-α (DAGLA) as the only protein whose palmitoylation consistently reflected both general Morris water maze experience and learning the hidden platform location across both paradigms. Together, these findings establish S-palmitoylation as a dynamic regulator of experience-dependent hippocampal plasticity.

## Introduction

The formation and stabilization of memory traces involve temporally distinct processes, ranging from rapid changes occurring within minutes to long-lasting adaptations that persist for days (Alberini, 2011; Tetzlaff et al., 2013; Ziegler et al., 2015). Early phases rely on activity-dependent signaling cascades and post-translational modifications that rapidly alter neuronal function. In contrast, later phases require the synthesis of new proteins, structural remodeling, changes in cellular organization, and sustained alterations in protein composition that support memory consolidation (Basu & Siegelbaum, 2015; Huang, 1998; Reymann & Frey, 2007).

Post-translational modifications (PTMs) are key regulators of these processes, enabling rapid and reversible control over protein localization, stability, and function. Among them, S-acylation, commonly referred to as S-palmitoylation (S-PALM) or simply palmitoylation due to predominant incorporation of 16-carbon palmitic acid moiety (Petropavlovskiy et al., 2021; Ramzan et al., 2023), is uniquely suited to act as a dynamic molecular switch. Through the reversible attachment of non-polar lipid chains, S-PALM regulates membrane association and trafficking of numerous proteins (Buszka et al., 2023; Chamberlain & Shipston, 2015; Fukata & Fukata, 2010; Matt et al., 2019). The function of S-PALM has been broadly characterized for scaffolding proteins PSD-95 and gephyrin, as well as ionotropic receptors such as AMPARs, NMDARs, and GABA receptors, which mediate fast synaptic transmission [reviewed in (Buszka et al., 2023; Zaręba-Kozioł et al., 2018)]. Several proteins regulated by palmitoylation, i.e., PSD-95, AKAP79/150, Arc, and cyclin Y, have been implicated in long-term potentiation and depression (Barylko et al., 2018; Chowdhury & Hell, 2019; Keith et al., 2012; Li et al., 2023; Seo et al., 2022; Shen et al., 2022). Experimental studies further support a functional role of S-PALM in behavior: inhibition of palmitoylation impairs spatial and contextual memory, while genetic manipulation of enzymes controlling palmitoylation dynamics alters synaptic strength and memory performance (Li et al., 2023; Milnerwood et al., 2013; Sapir et al., 2019; Urrego-Morales et al., 2023). Moreover, learning induces rapid changes in the palmitoylation state of numerous hippocampal proteins, including those involved in synaptic signaling, cytoskeletal organization, and metabolism (Nasseri et al., 2022).

Although the majority of palmitoyltransferases are localized in the Golgi apparatus and endoplasmic reticulum, several members of this family, as well as certain thioesterases, are present at or near synapses (Brigidi et al., 2015; Kim et al., 2008; Noritake et al., 2009; Pytyś et al., 2025; Thomas et al., 2012), enabling rapid and activity-dependent remodeling of the synaptic palmitoylome. Indeed, changes in protein palmitoylation can occur within minutes following synaptic stimulation (Brigidi et al., 2015; Pytyś et al., 2025). However, despite growing evidence supporting the role of S-PALM in functional and structural plasticity (reviewed in: Ji & Skup, 2021), hippocampal palmitoylome remodeling during learning and memory formation remains poorly understood. In particular, the dynamic regulation of the rat hippocampal palmitoylome during learning remains largely unknown.

Here, we sought to characterize the hippocampal palmitoylome following short and long hippocampus-dependent spatial learning protocols in the Morris water maze. To this end, palmitoylated proteins were analyzed using acyl-biotin exchange followed by isobaric labeling using Tandem Mass Tags and mass spectrometry (ABE-TMT LC-MS/MS). We identified distinct sets of neuronal proteins undergoing dynamic palmitoylation across learning phases and mapped specific cysteine residues targeted by this modification. These findings provide a proteome-wide view of S-palmitoylation dynamics *in vivo* and highlight its role in neuronal plasticity during learning and memory formation.

## Results

### Spatial learning performance in short- and long-term paradigms

To assess learning-related changes in protein S-palmitoylation, rats underwent hippocampus-dependent spatial training in the Morris water maze, where they learned to locate a hidden platform in a water-filled pool using distal visual cues for spatial orientation (**Figure 1A, C**, upper panels). The initial improvement in performance observed during the first trials of training primarily reflects short-term/working memory processes before spatial information is consolidated into long-term reference memory (Baldi et al., 2005). Therefore, one group of animals underwent short-term training consisting of 15 acquisition trials followed by a probe trial (STT, *n* = 8), whereas a second group underwent long-term training consisting of four training sessions (four trials per session) performed over four consecutive days, followed by a probe trial 24 h after the final session (LTT, *n* = 8) (**Figure 1A**; see Methods for details). To distinguish molecular changes specifically associated with learning the hidden platform location from those induced by general task experience, yoked control groups (n = 8 per group) underwent the same handling and swimming procedures but without a hidden platform. Although yoked animals experienced the same testing context and could acquire experience unrelated to platform localization, the removable distal visual cues were intentionally varied, preventing them from forming a stable memory of the hidden platform location (**Figure 1A**; see Methods for details).

**Figure 1.**
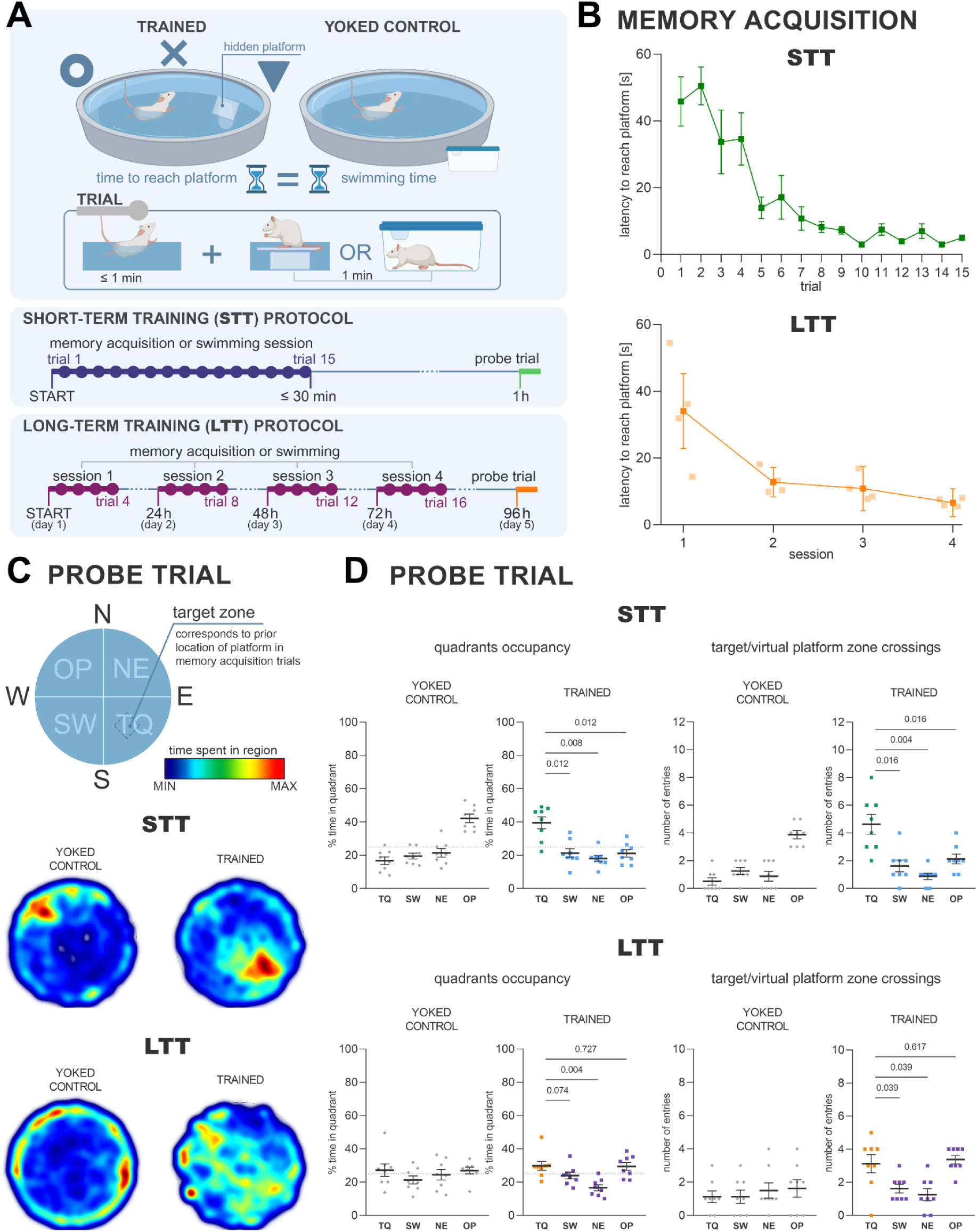
Overview of short- and long-term spatial learning paradigms. (**A**) Schematic of the Morris water maze experimental design. Animals were assigned to pairs consisting of a ‘trained’ and a ‘yoked’ control individual. Trained rats underwent spatial memory training to locate a hidden platform. Yoked rats swam in the pool without a hidden platform, while the removable distal visual cues were varied to prevent stable spatial learning. In the short-term training (STT), animals completed a single intensive session of memory acquisition (trained) or swimming (yoked) consisting of 15 trials (each trial included up to 1 min of swimming followed by 1 min of rest on the platform [trained] or in a cage adjacent to the pool [yoked]). This session was followed by a 1-min probe trial conducted 1 h after the start of the experiment. In the long-term training (LTT), animals underwent training or swimming sessions over 4 consecutive days (4 trials per day), followed by a 1-min probe trial on day 5. For details, see Methods. (**B**) Mean latency of trained animals to locate the platform during the STT (top, shown in green) and LTT (bottom, shown in saturated orange) protocols (n = 8 per group). Additionally, mean latencies for each trial within each LTT session are shown in lighter shade. In the STT group, escape latency significantly decreased across trials (Friedman test, χ²(14) = 70.03, *p* < 0.0001). Planned post hoc Wilcoxon signed-rank tests with Holm correction demonstrated significant reductions in escape latency between Trials 1-4 and Trials 5-8 (*p* = 0.0117), as well as between Trials 5-8 and Trials 9-12 (*p* = 0.0234). In contrast, escape latency did not differ significantly between Trials 9-12 and Trials 13-15 (*p* = 0.1484), indicating that animals had reached asymptotic performance and that no further statistically significant improvement in spatial learning occurred during the final phase of training. In the LTT group, escape latency significantly differed across training sessions (Friedman test, χ²(3) = 19.95, *p* < 0.001). Post hoc Wilcoxon signed-rank tests with Holm correction showed a significant reduction in latency from Session 1 to Session 2 (*p* = 0.023) and from Session 3 to Session 4 (*p* = 0.023), but not between Session 2 and Session 3 (*p* = 0.641), indicating a transient plateau in spatial learning. (**C**) Heatmaps showing the mean occupancy in the pool area by trained and yoked animals during probe trials in the STT and LTT groups. Warmer colors indicate longer occupancy. Probe trials were conducted in the absence of the platform. The top panel marks the quadrants of the pool: target quadrant [TQ], southwest [SW], opposite [OP], northeast [NE] and the target zone corresponding to the prior location of the hidden platform during the acquisition phase of the STT- and LTT-trained animals. (**D**) Behavioral performance measures in the probe trials for STT and LTT groups. At the STT probe trial, trained rats spent significantly more time in the target quadrant (39.5 ± 3.6%) than in the SW (*p* = 0.0117), NE (*p* = 0.0078), and opposite (OP; *p* = 0.0117) quadrants (one-tailed Wilcoxon tests). They also crossed the former platform position (target zone) more frequently (4.6 ± 0.7 times) than the corresponding virtual platform positions in the SW (*p* = 0.0156), NE (*p* = 0.0039), and OP (*p* = 0.0156) quadrants (one-tailed Wilcoxon tests). At the LTT probe trial, trained rats spent more time in the target quadrant (29.8 ± 2.7%) than in the NE quadrant (*p* = 0.0039; one-tailed Wilcoxon test), whereas the differences compared with the SW (*p* = 0.074) and opposite (OP; *p* = 0.727) quadrants were not significant. Likewise, rats crossed the former platform position (target zone; 3.1 ± 0.6 times) more frequently than the corresponding virtual platform positions in the SW (*p* = 0.039) and NE (*p* = 0.039) quadrants, but not in the OP quadrant (*p* = 0.617; one-tailed Wilcoxon tests). Within-group statistical analyses of quadrant occupancy and target-zone crossings were performed only for trained animals, as yoked animals were not trained to locate a hidden platform and therefore had no defined target location. The relatively higher occupancy of the opposite quadrant (OP), particularly in yoked animals, reflects the fixed release position at the beginning of the probe trial rather than goal-directed search. Data are presented as mean ± SEM; each point represents one animal (n = 8 per group). Exact adjusted P values are shown above the horizontal bars. All post hoc p values were adjusted using the Holm procedure.

STT-trained rats showed a rapid reduction in escape latency across consecutive training trials, with significant decreases during the early phase of acquisition, whereas performance stabilized between trials 9-12 and 13-15 (**Figure 1B**, upper panel). During the probe trial, STT-trained rats spent significantly more time in the target quadrant and crossed the former platform location more frequently than the corresponding virtual platform locations in the remaining quadrants, indicating successful spatial learning. LTT-trained rats also exhibited a progressive reduction in escape latency over four training sessions. The largest decrease in latency occurred between sessions 1 and 2, followed by a further reduction between sessions 3 and 4, whereas no significant change was detected between sessions 2 and 3 (**Figure 1B**, lower panel).

Heatmaps of spatial occupancy during the probe trials showed concentrated searching around the target zone in STT-trained rats, more dispersed search behavior in LTT-trained rats, and predominantly thigmotactic (wall-following) behavior in yoked controls (**Figure 1C**). Quantitative analysis confirmed these observations (**Figure 1D**). In the STT group, rats spent significantly more time in the target quadrant and crossed the target zone more frequently than the corresponding locations in the other quadrants, whereas yoked controls showed no preference for either the target quadrant or target zone (**Figure 1D**, upper panels). In the LTT group, spatial learning was less pronounced. Trained rats spent more time in the target quadrant than in the NE quadrant, but not the SW or opposite quadrants (**Figure 1D**, lower panels). Likewise, target-zone crossings exceeded those in the SW and NE quadrants, but not in the opposite quadrant. LTT yoked controls again showed no preference for the target quadrant or target zone (**Figure 1D**, lower panels). An additional analysis of search strategies (**Table 1**, **Supplementary Figure 1**), classified according to the algorithm of (Garthe et al., 2014), revealed a progressive shift from non-spatial to spatially guided navigation in both training protocols. During the initial trials, rats predominantly displayed thigmotaxis and other non-specific search strategies. With continued training, these behaviors were gradually replaced by directed search, corrected search, and ultimately direct path strategies, indicating the acquisition of an accurate spatial representation of the platform location. This transition occurred progressively across the four training days in the LTT group, whereas STT animals exhibited a comparable shift within a single intensive training session. Thus, despite the markedly different temporal organization of training, both protocols resulted in the adoption of increasingly efficient hippocampus-dependent spatial search strategies.

Together, these findings indicate that both training protocols successfully supported the acquisition of spatial navigation strategies. However, despite this convergence in search strategy, the subsequent probe trial revealed stronger spatial memory expression after STT than after LTT.

**Supplementary Figure 1.**
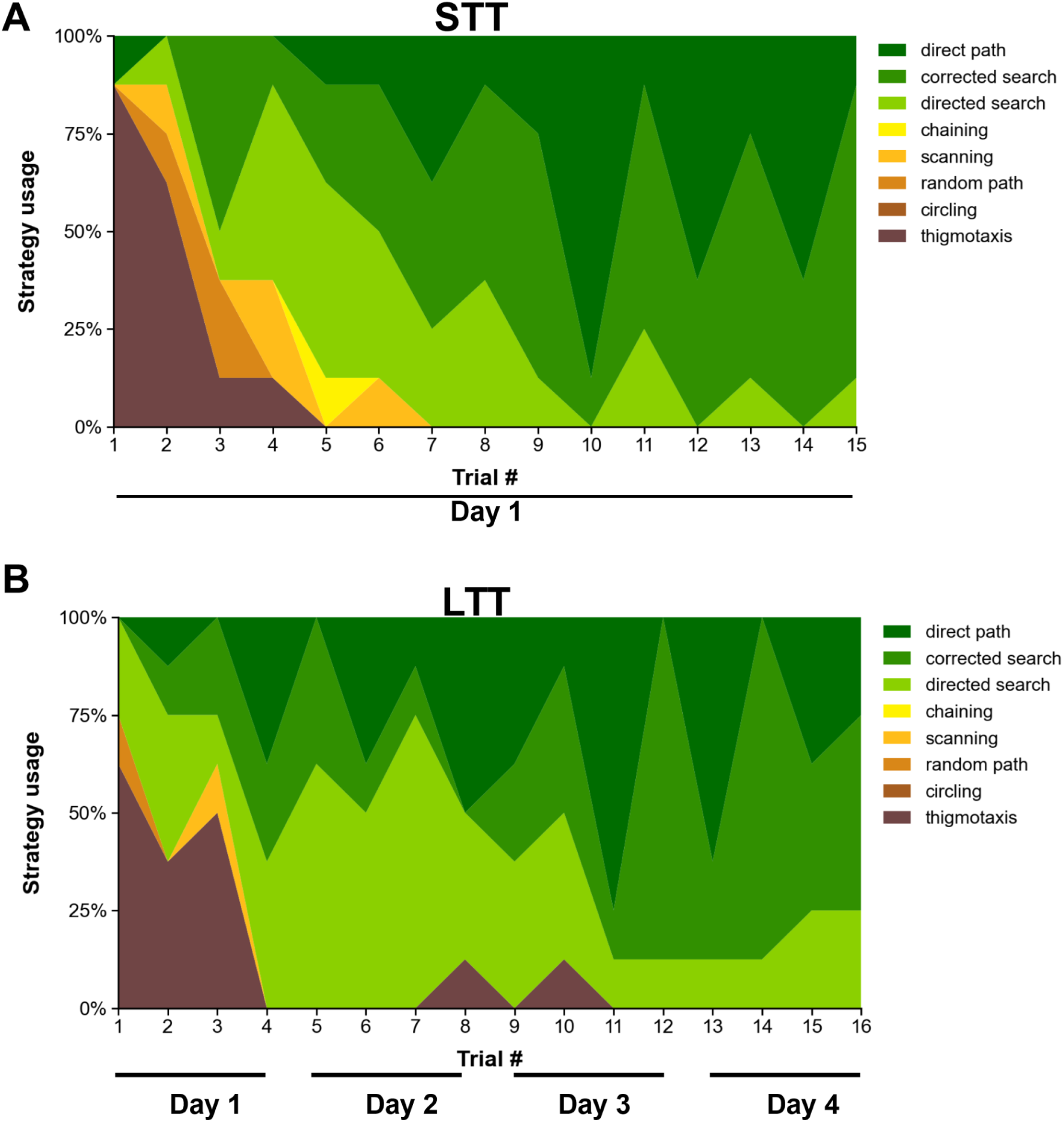
Contribution of search strategies in short-term training (**A**, STT) and long-term training (**B**, LTT) in the Morris water maze, determined by algorithm-based classification. Individual swim paths were classified using the Rtrack algorithm into predefined spatial search strategies. Colored areas indicate the percentage of animals using each strategy in each acquisition trial. Seven strategies of potential eight, listed and color coded, were used.

### Identification of differentially palmitoylated hippocampal proteins

To investigate training-related changes in the hippocampal palmitoylome, STT- and LTT-trained rats as well as respective yoked animals were sacrificed immediately following the probe trial and their hippocampi were rapidly extracted and snap-frozen. Tissues were collected immediately after the probe trial in both groups - 1 h after training onset for STT group and on day 5, 96 h after training onset, in the LTT group. In addition, cage control (naive) animals from the same cohort were sacrificed. Their tissue served to establish the baseline protein palmitoylation levels in animals not exposed to the experimental procedures. Although eight animals were trained in each group for the spatial learning experiments, six animals per group were randomly selected for mass spectrometry analysis because of technical limitations on the number of samples that could be analysed simultaneously within a single experimental run. The acyl-biotin exchange (ABE) method was used to tag and enrich palmitoylated proteins extracted from the rat hippocampi. Samples were subsequently labeled with tandem mass tags (TMT) and analyzed by LC-MS/MS (**Figure 2A**; see Methods section for details)

**Figure 2.**
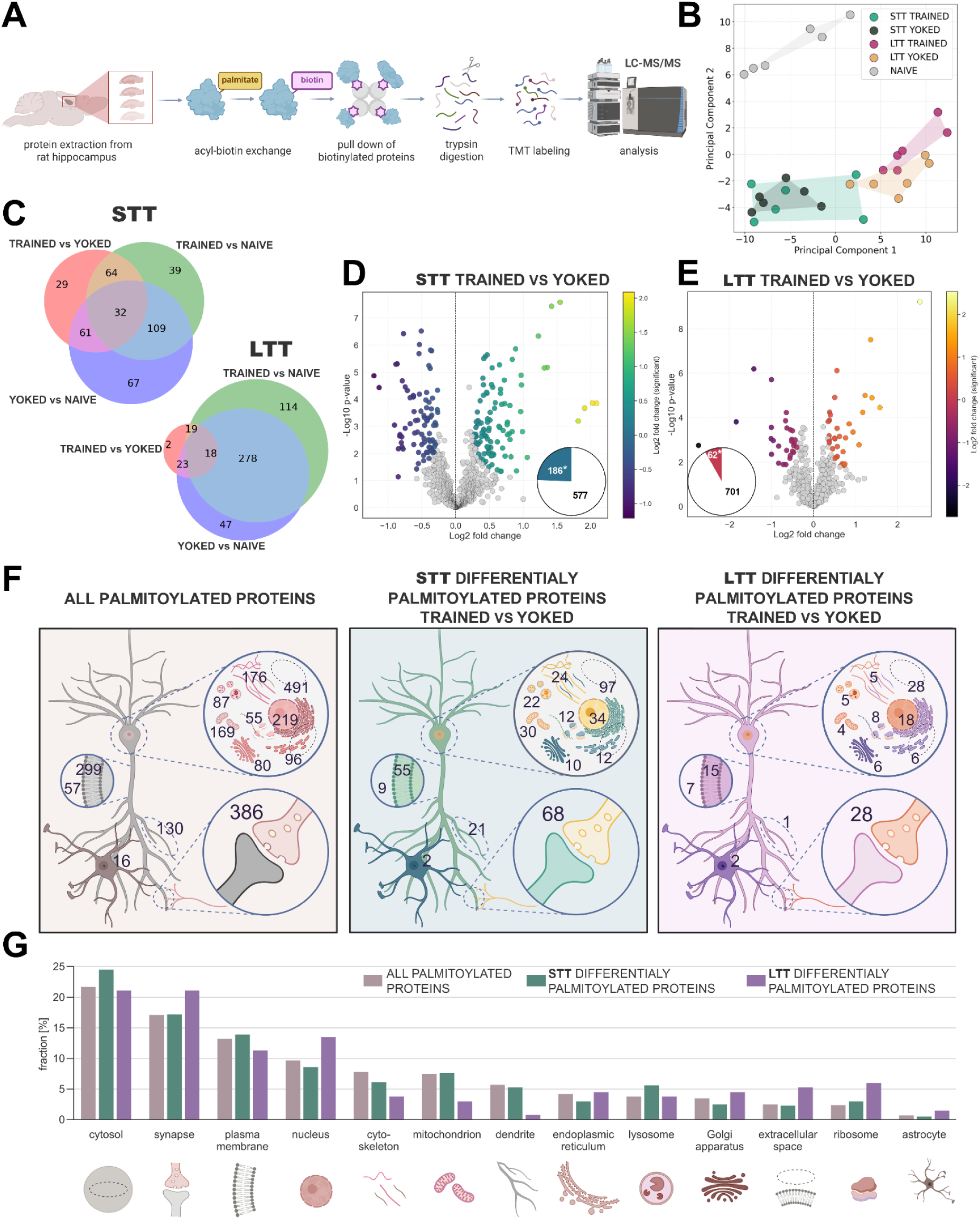
Remodeling of the hippocampal palmitoylome following spatial learning. **(A)** Diagram of the protocol for identification of palmitoylated proteins in rat hippocampal homogenates using acyl-biotin exchange assay coupled with Tandem Mass Tag labeling and liquid chromatography-tandem mass spectrometry. **(B)** Principal component analysis (PCA) of palmitoylated protein samples from naive animals, short-term (STT) and long-term (LTT) trained and yoked groups. Each point represents the palmitoylome profile of an individual animal (n = 6). **(C)** Venn diagrams showing the overlap between differentially palmitoylated proteins in the STT (left) and LTT (right) conditions. Numbers indicate proteins unique to or shared among the indicated groups. **(D-E)** Volcano plots comparing palmitoylation levels of individual proteins in STT (C) and LTT (D) groups versus their respective yoked controls. The x-axis represents the fold change in palmitoylation ([log_2_ intensity (trained - yoked values)], and the y-axis shows −log₁₀(p-value). A paired, two-sided Student’s *t*-test (S₀ = 0.1, FDR = 0.05) was used to identify proteins with significant changes in palmitoylation levels. Differentially palmitoylated proteins (DPPs) are shown in colour. Pie charts indicate the proportion of DPPs relative to the total palmitoylome for each training type. **(F)** Localization of all palmitoylated proteins (beige panel) and DPPs in STT and LTT groups (green and purple panels respectively), annotated using simplified UniProt Cellular Compartments descriptors (see Methods). Schematic representations show the distribution of proteins across various compartments, with numbers indicating the counts of palmitoylated proteins annotated to each compartment. **(G)** The bar graph summarizes the relative proportion of proteins assigned to each localization category, revealing a largely consistent representation of protein localization annotations across the compared sets of palmitoylated proteins. No significant differences were observed between the groups (Chi-square test: χ² = 15.43, df = 24, *p* = 0.90).

In total, we identified 5,260 proteins, of which 763 were palmitoylated (14.5%; **Table 2**; n = 6 animals per group). Significant changes in protein expression were detected in only 47 proteins (**Table 2**). This indicates that the observed alterations in the detection of palmitoylated proteins are not directly driven by changes in protein abundance. Additionally, in our analyses, the palmitome data were normalized to protein expression levels, accounting for potential effects of differential protein abundance on the observed palmitoylation changes. Principal component analysis (PCA) of the palmitoylome across all biological replicates revealed a clear separation between STT and LTT groups (including both trained and yoked animals), and from the naive control group, indicating time- and learning-dependent remodelling of the hippocampal palmitoylome (**Figure 2B**). We next compared differentially palmitoylated proteins (DPPs) for the experimental groups pairwise (**Figure 2C**). The greatest numbers of DPPs were identified when STT- and LTT-trained animals were compared with naive controls. Notably, the LTT-trained group showed the largest number of DPPs relative to naive animals, whereas only a small subset of proteins exhibited palmitoylation changes specifically associated with the acquisition of spatial memory for the hidden platform location, as revealed by the LTT-trained versus LTT-yoked comparison. Intermediate numbers of DPPs were observed in comparisons of both yoked groups versus naive animals. Comparisons between trained animals and their corresponding yoked controls yielded fewer DPPs, particularly in the LTT group. Altogether, these results indicate that general task experience, including environmental exposure, handling, physical exercise, and stress associated with the Morris water maze, induces pronounced changes in the hippocampal protein palmitoylome. In contrast, only a subset of these changes was specifically associated with learning the hidden platform location. Notably, a substantially larger pool of proteins underwent learning-specific palmitoylation changes immediately after STT than after LTT. In support of this, volcano plots illustrating changes in protein palmitoylation in STT- and LTT-trained groups relative to their yoked controls are shown in **Figure 2D-E** (see also **Table 3**). An analogous comparison between yoked animals and cage controls is shown in **Supplementary Figure 2**.

### Short- and long-term spatial learning differentially affects hippocampal palmitoylome

We next focused on comparisons between trained and yoked animals to isolate palmitoylation changes specifically associated with learning the hidden platform location. We asked what patterns of subcellular localizations could be ascribed to the trained vs. yoked DPPs identified in the STT and LTT groups. To this end, all identified S-palmitoylated proteins in hippocampal homogenates, as well as DPPs, were annotated using Subcellular Location [CC] terms from the UniProt database. More than 300 annotation terms were subsequently consolidated into 13 major annotation categories, including cellular and subcellular localizations such as cytosol, synapse, plasma membrane, nucleus, cytoskeleton, mitochondrion, dendrites, endoplasmic reticulum, lysosome, Golgi apparatus, ribosome, and extracellular space, as well as an astrocyte-associated category (**Figure 2F**; **Table 4**; the term dictionary is provided in the **Supplementary Information**). The bar graph in **Figure 2G** summarizes the relative distribution of proteins across these ascribed compartments, revealing comparable profiles between groups. Notably, synapses represented one of the most prominent locations for both the overall palmitoylated proteome and trained vs. yoked DPPs identified in STT and LTT groups. All in all, no significant differences in subcellular distribution were observed between overall identified palmitoylome, STT DPPs, and LTT DPPs (chi-square test: χ² = 15.43, df = 24, p = 0.90). These results suggest that learning the hidden platform location induces widespread changes in protein palmitoylation across multiple localization categories rather than in a site-selective manner, although synaptic proteins remain among the most prominently represented.

**Supplementary Figure 2.**
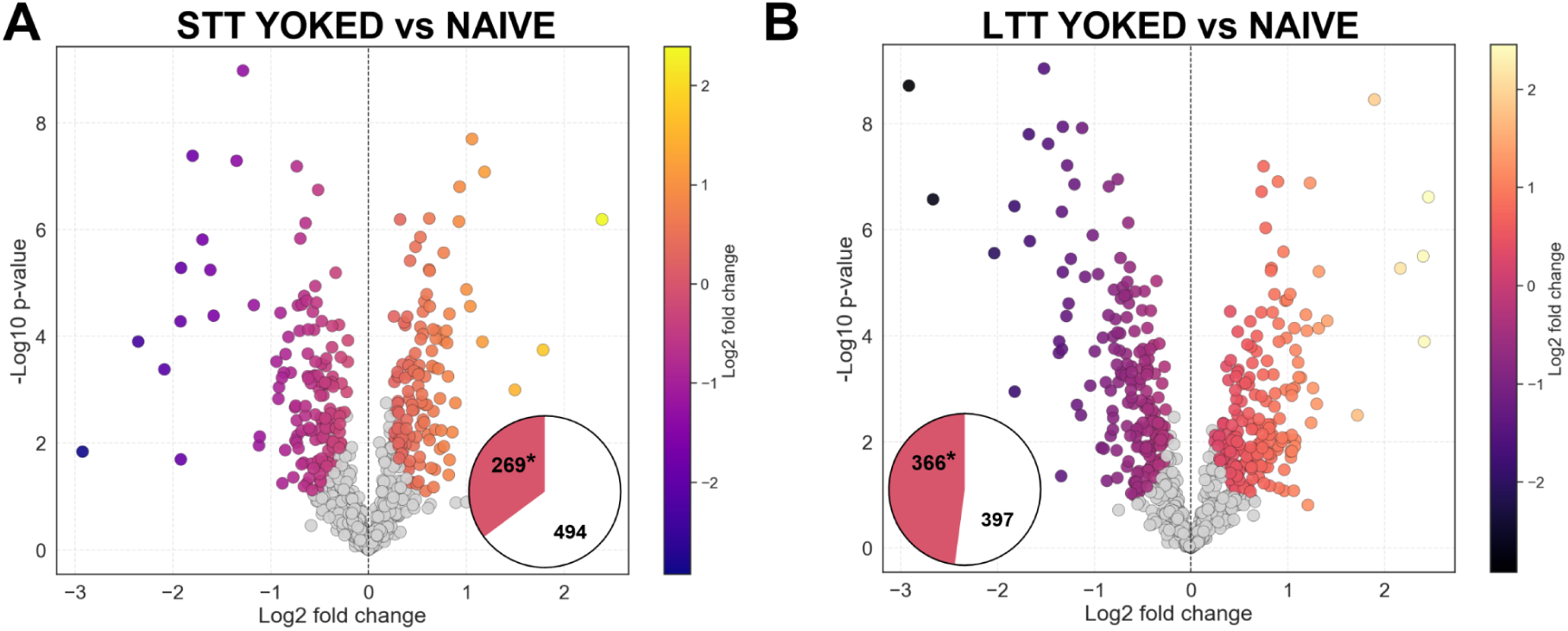
Remodeling of the hippocampal palmitoylome following environmental exposure independent of learning the hidden platform location. Volcano plots comparing palmitoylation levels of individual proteins in animals exposed to Morrie water maze in the absence of the platform (STT- (**A**) and LTT-yoked (**B**) groups versus cage controls (naive animals, n = 6 animals per group). The x-axis represents the fold change in palmitoylation (log_2_ intensity yoked - naive values), and the y-axis shows -log₁₀(p-value). A paired, two-sided Student’s *t*-test (S₀ = 0.1, FDR = 0.05) was used to identify proteins with significant changes in palmitoylation, and differentially palmitoylated proteins (DPPs) are shown in colours. Pie charts indicate the proportion of differentially palmitoylated proteins relative to the total palmitoylome for each training type.

To better visualize individual DPPs, hierarchical clustering based on log₂-transformed intensities is shown in **Figure 3**. The dendrograms illustrate clusters of proteins (rows) and samples (columns), grouping those with similar palmitoylation patterns. We identified clusters of proteins that were selectively hyper- or depalmitoylated following learning when compared with yoked controls (**Figure 3A-B**; **Table 3**). For example, in the STT group, in trained animals, syntaxin 7 (*Stx7*) and synaptic vesicle glycoprotein 2A (*Sv2a*) were hyperpalmitoylated relative to yoked controls. In contrast, proteins such as the G protein alpha subunit Gs-α (*Gnas*) and Ras-related protein Rab-2A (*Rab2a*) exhibited reduced palmitoylation (**Figure 3A**). In the LTT group, in trained animals, proteins such as neurexin-1 (*Nrxn1*) and neuronal cell adhesion molecule (*Nrcam*) were hyperpalmitoylated, whereas Ras-related protein Rab-2A (*Rab2a*) and sodium channel subunit beta-2 (*Scn2b*) showed decreased palmitoylation when compared with yoked controls (**Figure 3B**; **Table 3**). We further identified 24 proteins that exhibited differential palmitoylation in both the STT and LTT trained groups relative to their respective yoked controls (**Figure 3C**). Among these, 15 proteins exhibited consistent changes in the direction of changes to S-palmitoylation, whereas 9 proteins showed opposite regulation between the two different trainings (**Figure 3C**). These proteins can be ascribed to several functions including membrane trafficking and vesicle dynamics (Rab2a, Rab3b, Rab14, Igsf8, Dagla), lipid metabolism and membrane remodeling (Dagla, Acsf2), protein translation / RNA metabolism (Rps8, Rps29, Hnrnpa2b1, Sars1), cell signaling and neuronal plasticity (Iqsec2, Ndrg4), and metabolism / nucleotide metabolism (Prpsap2, Rnpep, Nipsnap1). Notably, these manually curated functional groups closely mirror the pathways identified by the unbiased enrichment analysis of the larger trained-versus-yoked DPP datasets, including membrane trafficking, synaptic organization, lipid metabolism, and protein translation. This convergence supports the biological relevance of the stringent hierarchical classification despite the limited number of candidate proteins. Altogether, these results suggest that short- and long-term learning affect relatively small and largely non-overlapping subsets of proteins in the hippocampus.

**Figure 3.**
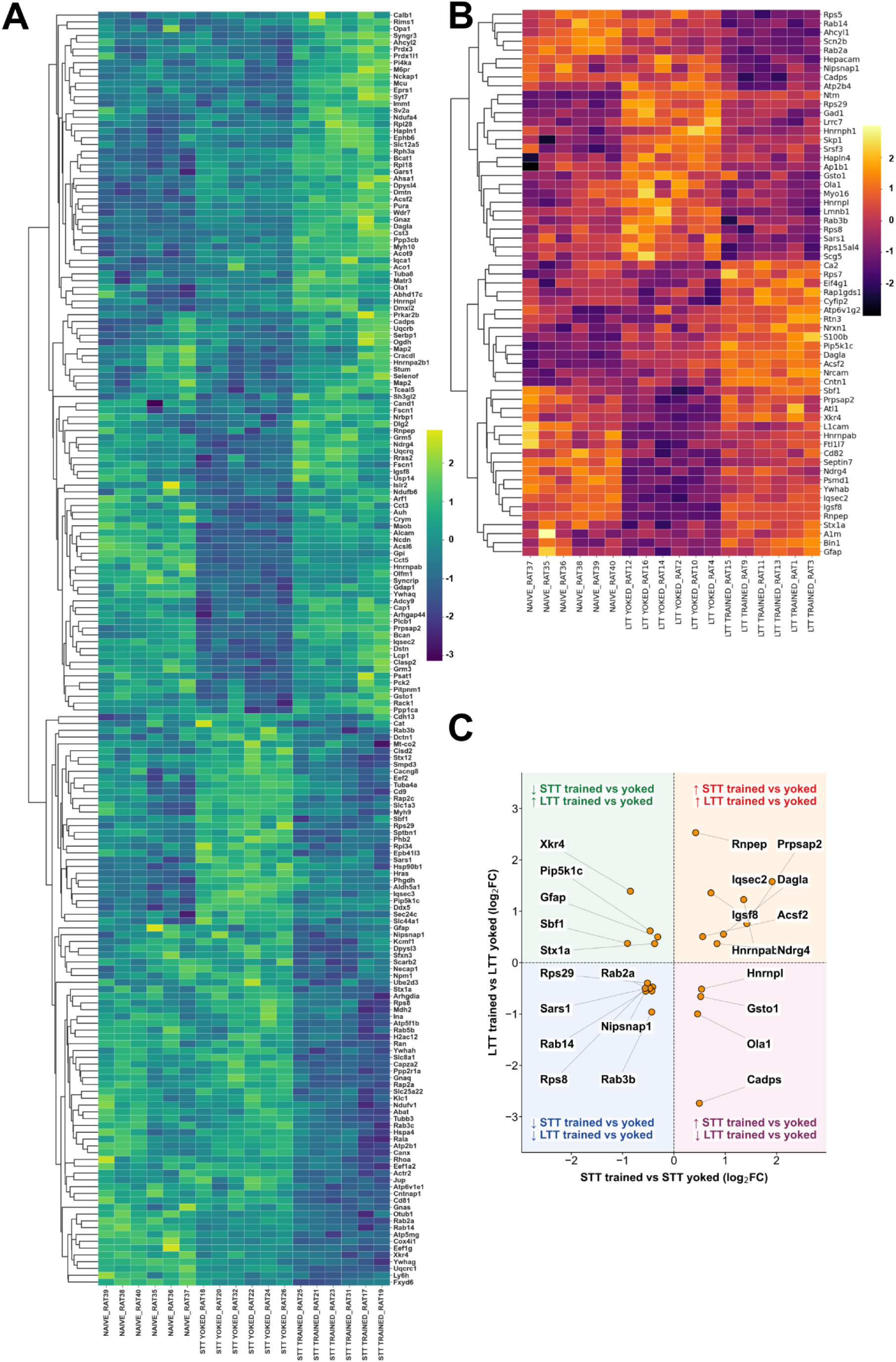
Clustering of differentially palmitoylated proteins in the hippocampus following spatial learning. Heatmaps of log_2_-transformed intensities for differentially palmitoylated proteins (DPPs) in short-term learning (STT) (**A**) and long-term learning (LTT) (**B**) groups, including trained and their respective yoked individuals as well as naive animals. Note that each protein is labeled using its corresponding gene name. Each row represents a protein of interest, and each column represents an individual animal sample. Colors indicate the magnitude of fold change. Dendrograms show hierarchical clustering of proteins (rows) and samples (columns), grouping those with similar fold-change patterns. **(C)** Proteins whose levels were significantly altered in both STT and LTT trained groups compared to respective yoked individuals (n = 24 common proteins; log_2_ intensities for trained - yoked groups). The quadrants distinguish proteins showing concordant or opposite directions of palmitoylation change between the two training paradigms.

To distinguish proteins responding to general Morris water maze (MWM) experience from those specifically associated with learning the hidden platform location, we performed a more stringent hierarchical classification of differentially palmitoylated proteins (DPPs). First, we identified proteins that exhibited significant, directionally consistent changes in palmitoylation across all four MWM-exposed groups (STT-trained, STT-yoked, LTT-trained, and LTT-yoked) relative to naive animals. We then determined which of these proteins additionally differed between trained and yoked animals, thereby identifying proteins whose general experience-dependent regulation was further modified by learning the hidden platform location (**Supplementary Figure 3**). This strategy allowed us to distinguish proteins associated with learning the platform location (trained-versus-yoked comparisons) from a more restricted subset of proteins that integrate both general behavioral experience and spatial learning. Using this approach, we identified 52 proteins that exhibited significant and directionally consistent palmitoylation changes in all four MWM-exposed groups relative to naive animals, including 24 proteins with increased and 28 with decreased palmitoylation. Among these, 14 proteins additionally differed between trained and yoked animals in at least one training paradigm, indicating that they respond to general MWM experience while also undergoing learning-specific regulation (**Supplementary Figure 3**). Notably, diacylglycerol lipase-alpha (DAGLA) was the only protein that fulfilled this stringent criterion in both the STT and LTT paradigms, identifying it as the strongest candidate linking experience-dependent palmitoylation with learning the hidden platform location.

**Supplementary Figure 3.**
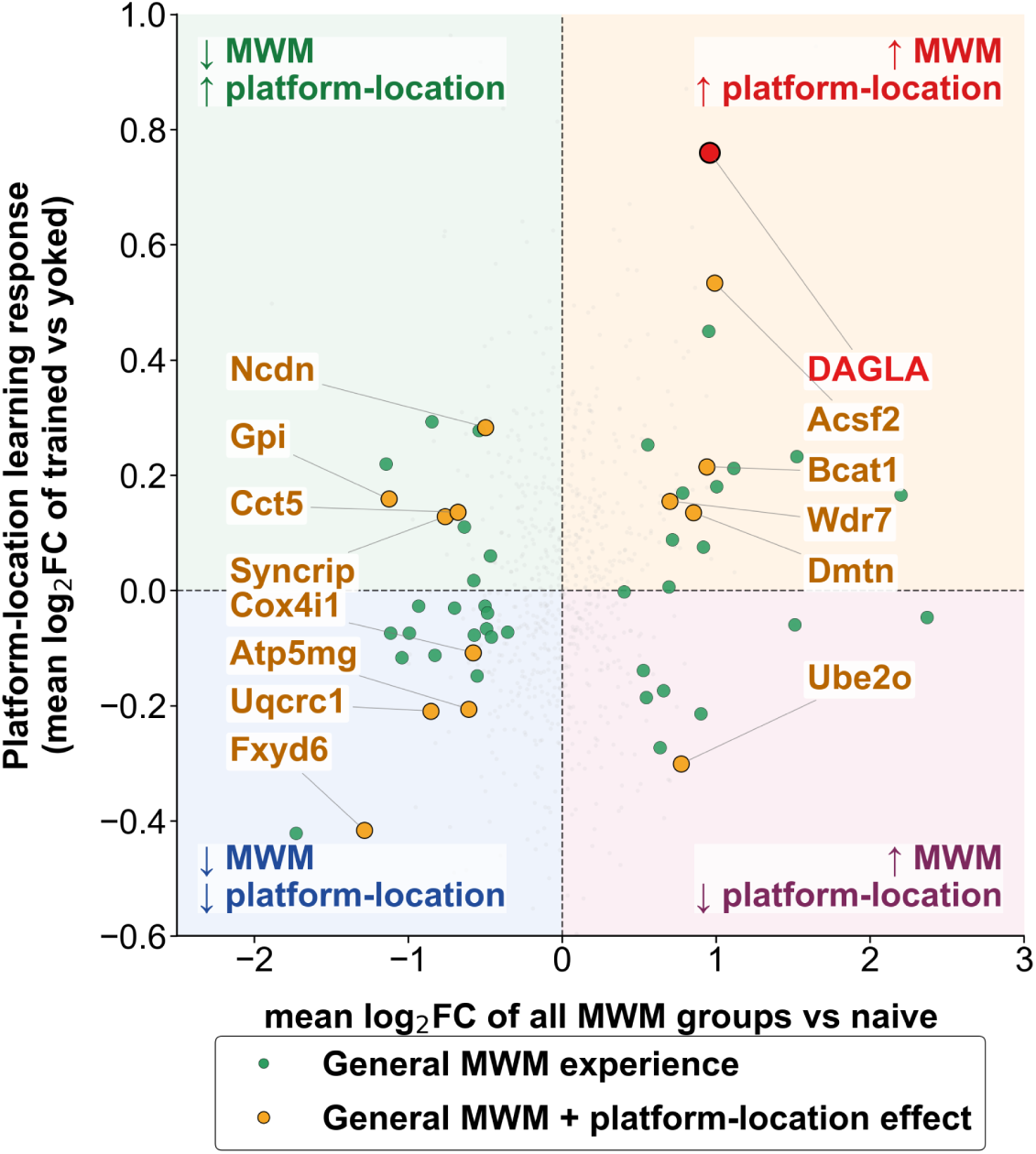
Hierarchical classification of experience- and learning-associated palmitoylation responses. Each point represents one protein. The x-axis shows the general MWM response, calculated as the mean log₂ fold change in palmitoylation across all four Morris water maze (MWM)-exposed groups (STT-trained, STT-yoked, LTT-trained, and LTT-yoked) relative to naive animals. The y-axis shows the platform-location learning response, calculated as the mean log₂ fold change between trained and the respective yoked animals across the STT and LTT paradigms. Green points denote proteins exhibiting significant and directionally consistent palmitoylation changes in all four MWM-exposed groups relative to naive animals (general MWM-responsive proteins). Orange points indicate proteins that additionally differed between trained and yoked animals in at least one training paradigm, identifying proteins whose general experience-dependent palmitoylation was further modified by learning the hidden platform location. Diacylglycerol lipase-α (DAGLA; red) was the only protein fulfilling this stringent criterion in both the STT and LTT paradigms. Quadrants indicate the direction of the general MWM-associated and platform-location-learning-associated palmitoylation responses.

To characterize the biological processes associated with DPPs identified in the trained-versus-yoked comparisons, we performed functional enrichment analysis (**Figure 4A-B**). Although the preceding hierarchical classification successfully prioritized proteins integrating both general MWM experience and platform-location learning, this stringent approach yielded only 14 candidate proteins, including a single protein (DAGLA) common to both the STT and LTT paradigms. Such small protein sets are unsuitable for robust pathway and network enrichment analyses. Therefore, to obtain a systems-level view of the biological processes associated with learning-dependent palmitoylation, subsequent functional enrichment analyses were performed using the complete DPP datasets from the STT and LTT trained-versus-yoked comparisons.

**Figure 4.**
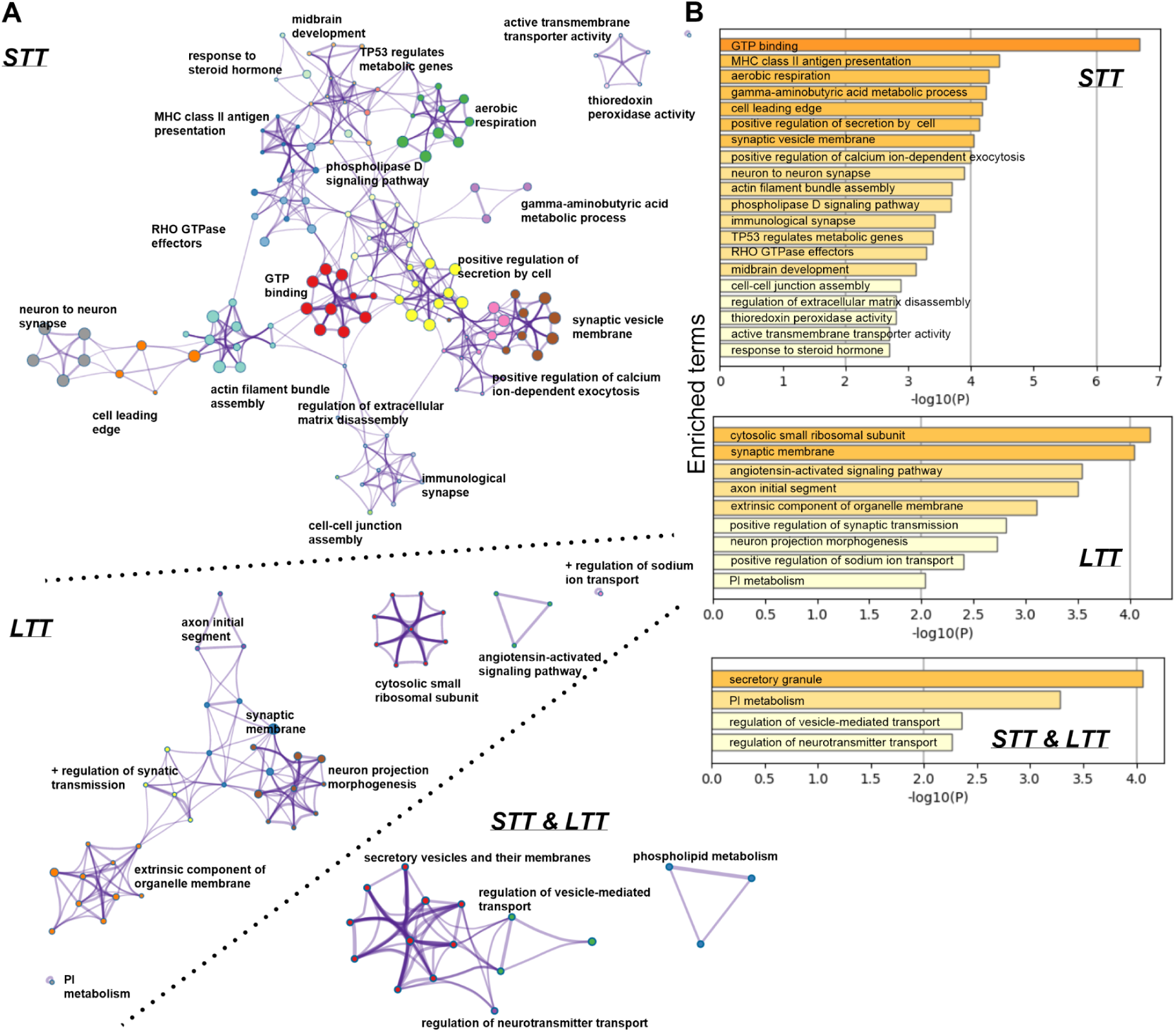
Functional enrichment analysis of differentially palmitoylated proteins in the hippocampus following the STT and LTT spatial learning protocols. **(A)** Graphical network representation of enriched terms associated with differentially palmitoylated proteins (DPPs) in the STT (upper network) and LTT (middle network) groups. An additional comparison of common DPPs identified in both STT and LTT groups is also shown (see Methods for details). A total of 186, 62, and 24 genes encoding palmitoylated proteins were analyzed, respectively. The input background database consisted of 5,087 genes encoding proteins and was used for enrichment calculations. Each term is represented by a node, the size of which is proportional to the number of genes assigned to that term, while node color indicates cluster identity (i.e., nodes of the same color belong to the same cluster). Terms with a similarity score > 0.3 are connected by edges, with edge thickness proportional to the similarity score. **(B)** Bar graphs showing enriched biological processes in the STT and LTT groups, as well as processes common to DPPs identified in both STT and LTT groups. Terms are colored according to p-value and hierarchically ordered based on kappa statistical similarities among their gene memberships.

In the STT group, pathway and process network analysis revealed 39 densely interconnected clusters corresponding to diverse biological processes (**Table 5**, **Figure 4A** top right, only top 20 clusters are shown for clarity). Enriched terms included synaptic function (e.g., synaptic vesicle membrane, neuron-to-neuron synapse, regulation of calcium ion-dependent exocytosis), signaling pathways involving GTP-binding proteins and Rho GTPases, actin cytoskeleton organization, and phospholipase D signaling. Additionally, processes related to mitochondrial function and metabolism (e.g., aerobic respiration, thioredoxin peroxidase activity), as well as gamma-aminobutyric acid (GABA) metabolic processes, were significantly enriched. The bar graph representation confirmed strong enrichment of terms such as GTP binding, synaptic vesicle membrane, and regulation of secretion, highlighting extensive and functionally diverse synaptic remodeling shortly after learning (**Figure 4B**, top panel). In contrast, the DPPs of the LTT group revealed a markedly smaller and more focused set of enriched processes. The enrichment networks were smaller and less interconnected, with dominant terms related to protein synthesis (e.g., cytosolic small ribosomal subunit), synaptic membrane organization, and regulation of synaptic transmission. Further functional enrichment terms revolved around neuronal structure and connectivity, including axon initial segment and neuron projection morphogenesis, as well as organelle membrane-associated components (**Figure 4A**, middle). The more compact nature of the enrichment terms ascribed to the DPPs of the LTT group is evident in the corresponding enrichment ranking (**Figure 4B**, middle), indicating a more restricted functional profile at later stages.

We additionally examined proteins that were identified as differentially palmitoylated in both STT and LTT conditions to pinpoint processes shared across spatial learning phases. The enrichment analysis of this common subset revealed clusters centered around secretory vesicles and vesicular membranes, regulation of vesicle-mediated transport, regulation of neurotransmitter transport, and phospholipid metabolism. Among the top enriched terms were secretory granule, phosphatidylinositol (PI) metabolism, regulation of vesicle-mediated transport, and regulation of neurotransmitter transport, suggesting persistent remodeling of membrane trafficking and lipid signaling pathways across both short- and long-term learning conditions (**Figure 4A-B**, bottom). Overall, these results demonstrate that early learning engages a broad spectrum of biological processes, including synaptic signaling, metabolism, and cytoskeletal dynamics, whereas long-term learning is associated with more selective enrichment of processes related to protein translation and synaptic organization. This shift suggests a transition from widespread cellular remodeling during initial learning to more targeted mechanisms supporting memory consolidation. Shared STT/LTT palmitoylation changes were enriched in vesicular trafficking and phospholipid regulatory pathways, indicating core mechanisms maintained across memory phases.

To further characterize the functional organization of DPPs, we performed protein-protein interaction (PPI) network analysis followed by Molecular Complex Detection MCODE clustering (**Figure 5A-B**). PPI analysis reconstructs and interprets how proteins physically or functionally interact inside a cell and can be built from both experimental/literature evidence and computational or database-predicted interactions. The resulting network includes proteins that physically interact with at least one other protein from the input list. For networks containing between 3 and 500 proteins, the MCODE algorithm was applied to identify densely connected clusters. The clusters identified by MCODE are shown in color. For the DPPs of the STT condition, the PPI network revealed a dense and highly interconnected structure with multiple distinct functional clusters. Prominent modules included proteins associated with mitochondrial respiration, ribosomal proteins and translation machinery, as well as signaling pathways involving G proteins and small GTPases. Additional clusters were linked to synaptic vesicle exocytosis and neurotransmitter release and non-proteinogenic amino acid metabolism including GABA metabolism (**Figure 5A**). These data are consistent with the results of the functional enrichment analysis. Analogous to the functional enrichment, the DPPs of the LTT group exhibited a markedly sparser PPI network with fewer and less interconnected clusters. The most prominent module was associated with ribosomal proteins and translation, suggesting sustained regulation of protein synthesis at later stages. Smaller clusters were related to synaptic adhesion and signaling, as well as vesicle trafficking and membrane-associated processes (**Figure 5B**). Next, we performed quantitative KEGG pathway over-representation analysis of DPPs in the STT and LTT groups. Key KEGG pathways included those related to neurodegenerative diseases, long-term potentiation and depression, and glutamatergic synapse in the STT group, as well as synaptic vesicle cycle and cell adhesion molecules in the LTT group (**Supplementary Figure 4**, **Table 6**). We then visualized DPPs from the STT and LTT groups based on their involvement in synaptic potentiation and other synaptic processes within KEGG pathways using the KEGG Mapper tool (https://tomaszwojtowicz-coder.github.io/KEGG/index.html). Overall, this analysis suggests that early learning may be associated with the engagement of multiple cellular systems, whereas later stages reflect more targeted regulation of synaptic and translational machinery during memory consolidation.

**Figure 5.**
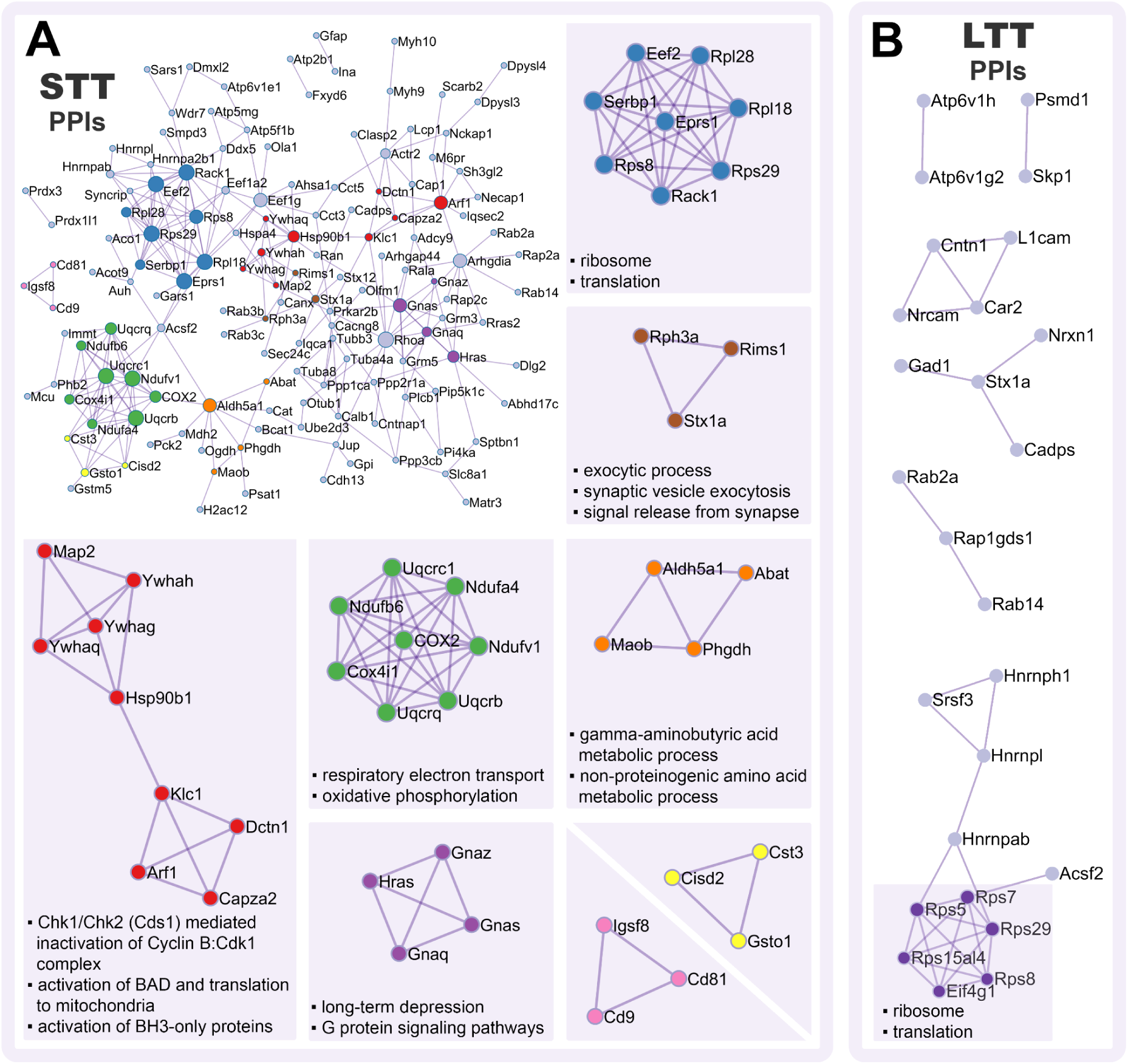
Protein-protein interaction enrichment analysis of differentially palmitoylated proteins. Protein-protein interaction (PPI) networks of DPPs identified in trained versus yoked comparisons were constructed using Metascape and visualized with Cytoscape, for STT **(A)** and LTT **(B)** groups. Only experimentally validated physical interactions from STRING and BioGRID were included. The resulting networks display proteins that physically interact with at least one other protein in the dataset.

**Supplementary Figure 4.**
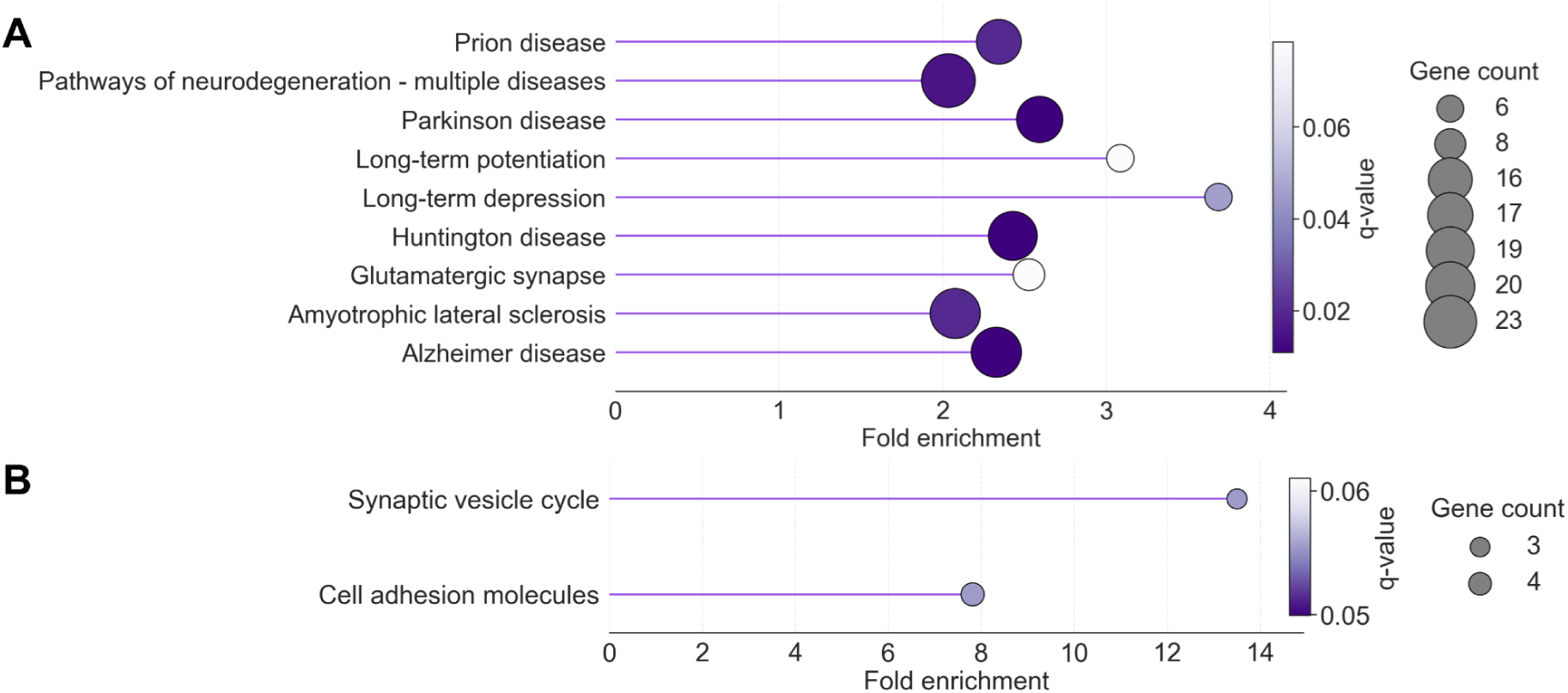
KEGG pathway analysis of differentially palmitoylated proteins. Quantitative KEGG pathway over-representation analysis (clusterProfiler, R) of differentially palmitoylated proteins in the STT group (**A**) and LTT group (**B**). Dot size indicates gene count, color represents FDR-adjusted q-values, and fold enrichment is shown on the x-axis.

A previous study in mice identified proteins that were differentially palmitoylated at 1 h in response to contextual fear-conditioning (Nasseri et al., 2022). We compared these proteins with homologous proteins identified in rats after STT in the present study. We were able to match 115 of 121 and 186 of 186 homologous proteins, respectively, with 23 proteins overlapping between the two datasets (**Supplementary Figure 5A**, **Table 7**). These proteins included cytosolic and cytoskeleton-related proteins, as well as postsynaptic signaling proteins and proteins associated with synaptic vesicles (i.e., including calcium voltage-gated channel auxiliary subunit gamma 8 and synaptic vesicle glycoprotein 2A, **Supplementary Figure 5B**). These findings suggest a partial conservation of palmitoylation dynamics across species and learning paradigms, with a subset of proteins consistently regulated following memory formation. The enrichment of cytosolic, cytoskeletal, and synaptic proteins among the common proteins indicates their diverse cellular localization. Moreover, palmitoylation-dependent remodeling of synaptic structure and function may represent a common mechanism among different species contributing significantly to early memory formation-related processes.

**Supplementary Figure 5.**
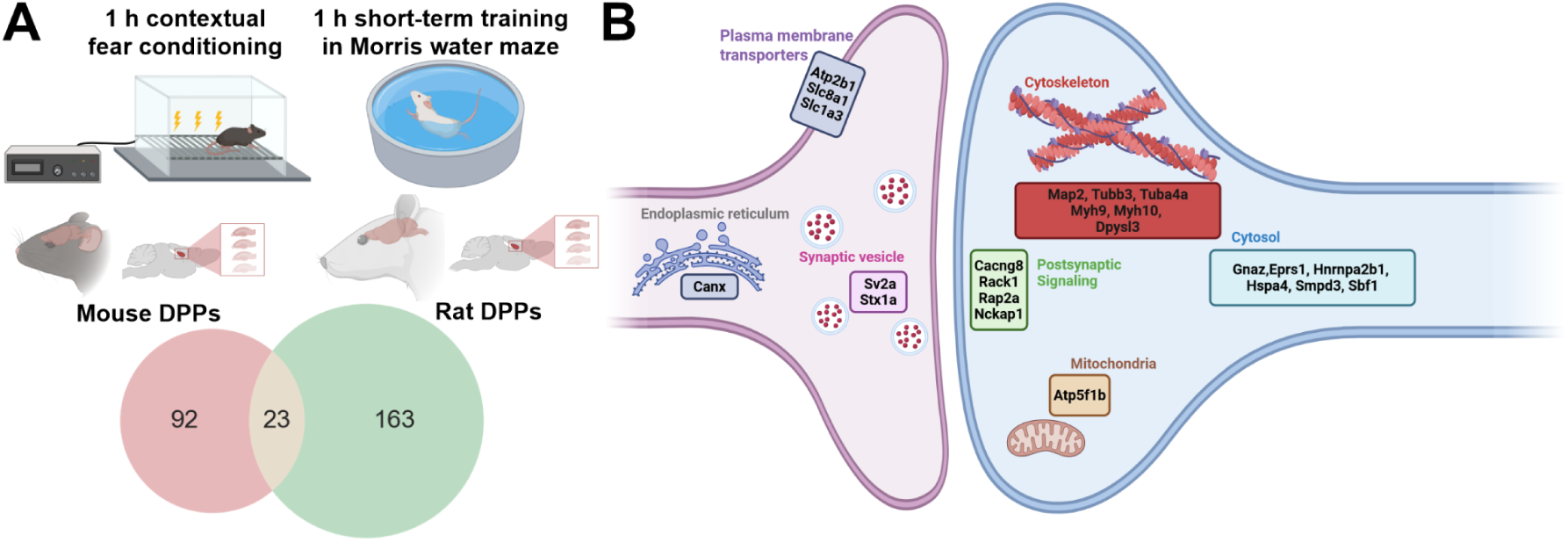
Comparison of differentially palmitoylated proteins 1 h after spatial learning in rats and contextual fear conditioning in mice. **(A)** Comparison of differentially palmitoylated proteins (DPPs) identified 1 h after contextual fear conditioning in mice (previous study by Nasseri et al., 2022) with those identified in rats after the STT protocol (this study). Gene symbols were converted to orthologous genes and verified for homology before comparison. **(B)** Schematic representation of DPPs common to both datasets

We next sought to determine the cysteine sites undergoing palmitoylation in proteins obtained from rat hippocampal tissue. To this end, proteins were extracted from homogenized hippocampi, and palmitoylated proteins were labeled using the ABE assay as previously described. However, unlike in the first experiment described above (**Figure 2A**), in which were proteins enriched by the ABE assay were subjected to tryptic digestion, proteins in this experiment were trypsin-digested prior to enrichment, and only peptides containing biotin-tagged cysteines were captured on resin, TMT-labeled, and analyzed by LC-MS/MS (**Figure 6A**, see Methods section for details). This approach allowed us to specifically identify individual palmitoylated peptides and locate the exact positions of palmitoylated cysteine residues in hippocampal proteins.

**Figure 6.**
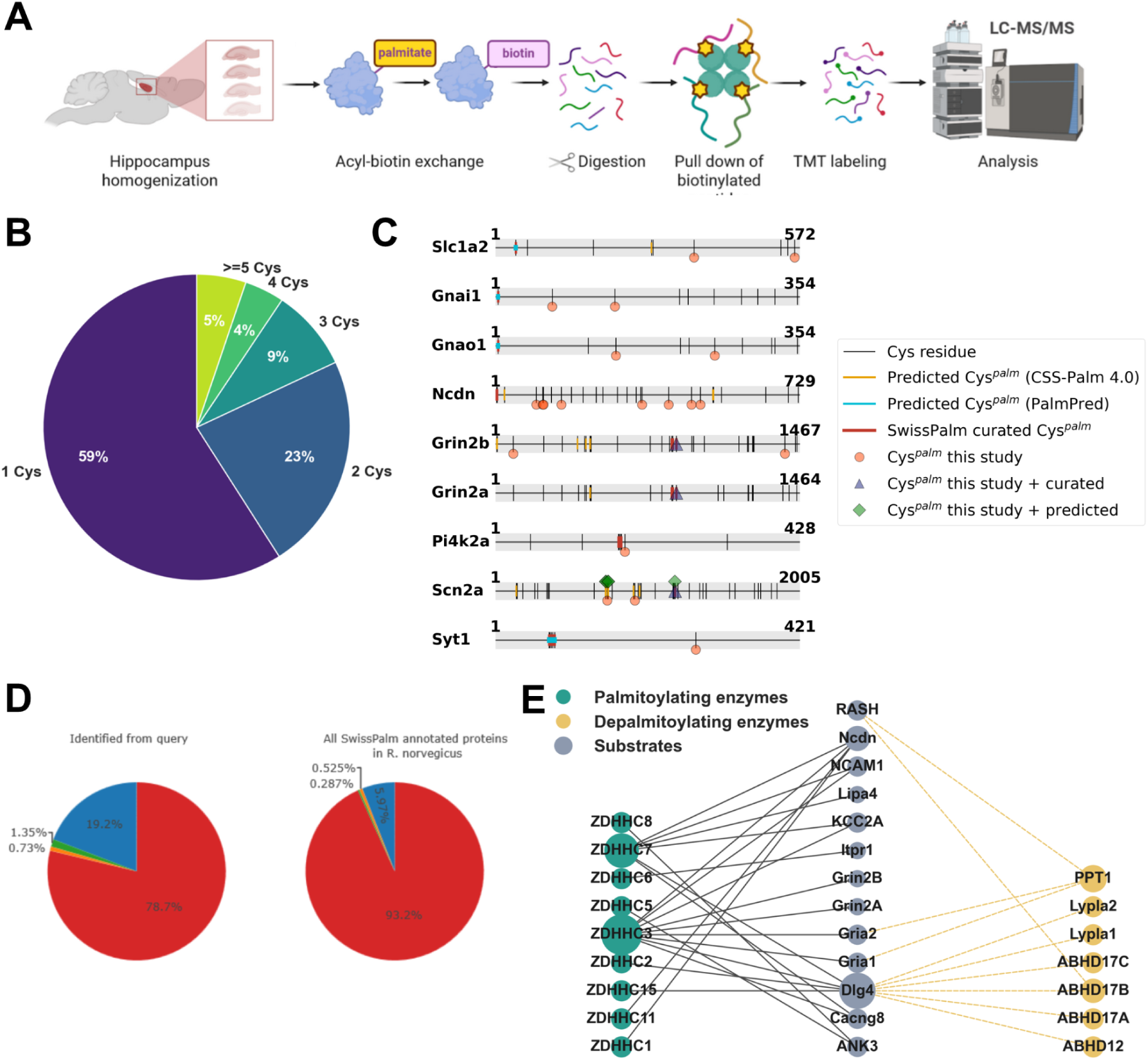
Identification of palmitoylated cysteine residues in rat hippocampal proteins. **(A)** Experimental workflow. Hippocampal homogenates from naive rats (n = 4) underwent palmitate substitution using the acyl-biotin exchange (ABE) method, followed by trypsin digestion. Palmitoylated peptides were enriched by pull-down and analyzed by LC-MS/MS. **(B)** Statistics of identified palmitoylated cysteines. Among the 1,482 proteins identified using this approach, most contained one or two palmitoylated cysteine residues. The full list of peptide sequences and identified cysteine residues is provided in Supplementary Table 1. **(C)** Rat protein cysteine palmitoylation mapping. Horizontal bars represent the full-length proteins, with amino acid positions indicated at each end. Black vertical lines mark cysteine residues in the protein sequences. Palmitoylation sites predicted with CSS-Palm 4.0 and PalmPred are shown alongside curated sites from the SwissPalm database in colours (amber, cyan and red, respectively). Circles indicate palmitoylation sites identified in this study (orange), including those unique to this study or overlapping with curated and/or predicted sites (other colours). **(D)** The relative fraction of proteins assigned to each cellular compartment depending on the number of identified palmitoylated cysteines. While the overall subcellular localization patterns were broadly conserved across groups, modest shifts in compartment representation were detected, particularly for ribosomal, Golgi apparatus, and lysosomal proteins. These distributions differed significantly among groups (Pearson chi-square test: χ² = 44.23, df = 22, p = 0.005). **(E)** SwissPalm database annotation of palmitoylated proteins identified during cysteine site identification (n = 1,482 proteins). Right panel: Fewer than 20% of proteins overlap with previously reported palmitoylomes in the SwissPalm database (blue), whereas more than 78% have not been previously reported as palmitoylated in *Rattus norvegicus* (red). Overlap with previously reported palmitoylomes and validation experiments (green), as well as proteins identified only in validation experiments (orange), is also shown. Left panel: Distribution of S-palmitoylated proteins identified in this study relative to all palmitoylated proteins annotated in SwissPalm (blue) and to the complete set of annotated rat proteins (>13,000; red). Colors as in the right panel. **(F)** Network representations of known palmitoyltransferases and depalmitoylating enzyme associations retrieved from SwissPalm that are relevant for proteins identified in B. Green and gold nodes indicate palmitoylating and depalmitoylating enzymes, respectively, and node size is proportional to the number of annotated enzyme-substrate interactions.

We identified a total of 2704 palmitoylated peptide sequences, which were ascribed to 1482 proteins (**Table 8**). Predominantly, we found 1-2 cysteine residues palmitoylated per protein (**Figure 6B**), and only a small fraction of proteins (5%) had ≥ 5 cysteine residues palmitoylated, with some proteins having 10-15 sites (clathrin, ankyrin-2, fatty acid synthase, or dynein, **Table 8**). We next compared our mapping results with the SwissPalm database [(Blanc et al., 2015), SwissPalm.org, last accessed January 1st 2026], which offers information about both experimentally confirmed palmitoylated cysteine positions in proteins of given species, as well as theoretical palmitoylation sites predicted with software such as CSS-Palm 4.0 (Ren et al., 2008) and PalmPred (Kumari et al., 2014). Currently, the database shares results of 2 previous studies in rats [(Edmonds et al., 2017; Kang et al., 2008)], and out of all rat proteins in the database (>13,000) less than 6% were reported to have been identified in S-PALM form. For 9 proteins in our dataset, we could find mapping information in the database. As shown in **Figure 6**, the majority of cysteine sites identified in this study are new, while some overlap with either curated cysteine sites or predicted sites (**Figure 6C**).

We next investigated whether the number of identified palmitoylated cysteine residues per protein was associated with specific localization annotations. Proteins were grouped according to their annotated localization and stratified by the number of palmitoylated cysteine residues (one, two to three, or four or more) (**Figure 6D**). Across all groups, cytosolic, synaptic, and plasma membrane proteins constituted the predominant fractions, indicating a broadly conserved distribution of palmitoylated proteins across localization categories. This pattern was consistent with the distribution observed in **Figure 2F-G**. However, modest but significant shifts in compartment representation were observed as the number of palmitoylated cysteines increased (Pearson chi-square test: χ² = 44.23, df = 22, p = 0.005). In particular, proteins containing four or more palmitoylated cysteines were underrepresented among ribosomal proteins, while they had increased representation amongst proteins ascribed to the Golgi apparatus. These findings suggest multiple palmitoylation sites may help proteins exert their function linked to membrane trafficking, organellar dynamics, and intracellular sorting pathways.

To place the identified palmitoylated proteins in the context of known enzymatic regulatory pathways, we next compared the 1,482 palmitoylated proteins identified in this study with proteins previously reported in the SwissPalm database. We found that fewer than 20% of proteins identified here overlapped with previously reported palmitoylomes in SwissPalm, whereas more than 78% had not been previously reported as palmitoylated in rats (**Figure 6D**, left panel). S-palmitoylated proteins identified in this study cover <6% of all palmitoylated proteins annotated in SwissPalm and to the complete set of annotated rat proteins (>13,000 proteins, **Figure 6D**, right panel). Altogether, this indicates that our dataset captures a distinct and previously underrepresented subset of the rat palmitoylome, substantially expanding the repertoire of known S-palmitoylated proteins and highlighting the limited coverage of palmitoylation events currently annotated in public databases.

We next searched the SwissPalm database for curated enzyme-substrate relationships corresponding to proteins identified in our dataset. This analysis retrieved 13 proteins for which palmitoyltransferases and depalmitoylating thioesterases had been experimentally validated in previous studies. The interaction network involving ZDHHC family palmitoyltransferases and depalmitoylating thioesterases, shown in **Figure 6F**, illustrates the potential complexity of the regulatory interactions that may underlie the broader set of differentially palmitoylated proteins identified in rat brain. For instance ZDHHC3 and ZDHHC7 emerged as prominent hubs with numerous annotated substrates, including PSD95 (Dlg4), glutamate receptor subunits, neurochondrin, and TARPγ8, consistent with their established roles in neuronal palmitoylation. In parallel, depalmitoylating enzymes such as PPT1, ABHD17 family members, LYPLA1/2, and ABHD12 were linked to overlapping subsets of substrates, highlighting the reversible nature of this modification. Several key synaptic proteins, including PSD95 and HRas, were connected to both palmitoylating and depalmitoylating enzymes, indicating dynamic bidirectional regulation.

## Discussion

In this study, we provide a comprehensive analysis of the hippocampal palmitoylome associated with spatial learning in the Morris water maze. By combining behavioral paradigms with quantitative proteomics, we demonstrate that protein S-palmitoylation in the hippocampus undergoes dynamic and phase-specific remodeling during memory formation. Our data reveal a temporal dissociation, with widespread and functionally diverse palmitoylation changes occurring shortly after acquisition of the hidden platform location (STT group), followed by a more restricted and functionally focused pattern after repeated training (LTT group). These findings support the idea that S-palmitoylation contributes to both rapid synaptic modulation during learning acquisition and longer-term processes associated with memory consolidation.

Previous research suggests that the Morris water maze robustly measures spatial learning when training is distributed across multiple days or when longer (≈2.0-2.5 h) intervals are used between sessions (Barrientos et al., 2016). In contrast, a compressed, one-day protocol (STT group) primarily reflects acquisition and within-session learning rather than long-term spatial reference memory (Baldi et al., 2005; Barrientos et al., 2016). Accordingly, the palmitoylation changes detected 1 h after the onset of the STT protocol most likely reflect molecular mechanisms supporting the acquisition of the hidden platform location and early memory formation. In this regard, one of the most prominent observations is the scale and diversity of palmitoylation changes during early learning. Approximately one quarter of the detected palmitoylome exhibited differential regulation at the 1-h time point (STT group), encompassing proteins involved in synaptic transmission, cytoskeletal dynamics, metabolism, and intracellular signaling. Our results corroborate a previous palmitoylome study performed in hippocampal homogenates 1 h after contextual fear conditioning in mice. Approximately 35% of palmitoylated proteins were differentially palmitoylated, exhibiting either increased or decreased palmitoylation, and were associated with synaptic function, metabolism, cytoskeletal organization, and signal transduction (Nasseri et al., 2022). This broad engagement of multiple functional systems is consistent with the notion that initial memory encoding requires coordinated remodeling across cellular compartments. The enrichment of proteins associated with synaptic vesicle cycling, calcium-dependent exocytosis, and small GTPase signaling suggests that S-palmitoylation may regulate presynaptic release probability as well as postsynaptic signaling cascades. In parallel, the identification of cytoskeletal and actin-related pathways supports a role for palmitoylation in structural plasticity, potentially by modulating protein localization and membrane association during spine remodeling.

In contrast, the palmitoylation landscape observed following the LTT protocol was markedly more selective. Because this paradigm combines repeated learning of the hidden platform location with prolonged exposure to the behavioral task, the reduced number of differentially palmitoylated proteins identified specifically in the trained-versus-yoked comparison suggests that relatively few palmitoylation events remain uniquely associated with learning the platform location after repeated training. Instead, the persistent changes observed relative to naive animals likely reflect a combination of memory maintenance and experience-dependent adaptations induced by repeated behavioral exposure. It should also be considered that the probe trials in the STT and LTT paradigms were performed after different retention intervals (1 h and 24 h, respectively). Consequently, the more robust memory expression observed in the STT group may reflect not only differences in the training paradigms but also the shorter interval between training and testing. Likewise, the distinct palmitoylation profiles observed after STT and LTT may represent different stages of memory processing, encompassing both differences in training history and the temporal evolution of molecular mechanisms underlying memory retention.

An important aspect of this study is the use of yoked controls to distinguish molecular changes associated with learning the hidden platform location from those induced by general task experience. However, yoked animals should not be regarded as “non-learning” controls. Although they could not acquire a stable memory of the hidden platform location, they repeatedly experienced the same testing context, including swimming, handling, and invariant features of the environment. Because the removable distal visual cues were intentionally varied between trials, yoked animals were not provided with a stable spatial framework required for learning the platform location, although they could still acquire familiarity with the general testing context. Moreover, unlike trained animals, yoked rats had no behavioral control over the termination of the aversive swimming experience, potentially resulting in differences in stress perception and predictability. Consequently, the trained-versus-yoked comparison should be interpreted as identifying molecular changes specifically associated with learning the hidden platform location, rather than the presence or absence of learning itself.

We found that 14.5% of the identified hippocampal proteins under basal conditions were palmitoylated. The strong representation of synaptic proteins across all conditions further supports the notion that synapses are a major site of palmitoylation-dependent regulation, consistent with the known localization of key palmitoyltransferases and thioesterases at synapses. This agrees with previous reports indicating that approximately 10% of the genome and up to 41% of synaptic genes encode palmitoylated proteins (Sanders et al., 2015). Despite marked functional remodeling, learning-related palmitoylation did not preferentially target proteins assigned to any particular subcellular compartment, as the overall subcellular distribution of DPPs remained unchanged. Importantly, comparison with the yoked control groups revealed that general task experience alone induced extensive remodeling of the hippocampal palmitoylome, whereas only a relatively small subset of proteins exhibited palmitoylation changes specifically associated with learning the hidden platform location. Thus, much of the observed palmitoylome remodeling reflects experience-dependent plasticity, while a distinct subset of proteins appears to be selectively regulated during spatial memory formation. By directly comparing trained animals with time-matched yoked controls, we were able to isolate task-specific effects, revealing that a distinct subset of palmitoylation changes is uniquely associated with memory formation rather than general environmental exposure. In addition, we detected expression of acyl-CoA synthetase long chain family proteins (ACSL1, ACSL3, ACSL4, ACSL5, ACSL6) and fatty acid synthase (FASN) in hippocampal homogenates. Importantly, in our previous analysis of the synaptoneurosomal palmitoylome, these same enzymes were also identified within the synaptic compartment (Pytyś et al., 2025). Together, these findings support a mechanistic model in which both activation of long-chain fatty acids to acyl-CoA by ACSL enzymes and *de novo* fatty acid synthesis mediated by FASN are locally available in hippocampal neurons, including at synapses. Such a spatial organization would enable tight coupling between lipid metabolism and S-palmitoylation cycles, providing a substrate supply that can directly regulate dynamic protein palmitoylation at or near synaptic sites. However, it remains unclear how the activity of these enzymes, which may contribute to a metabolic-posttranslational interface, is regulated during learning.

A prominent feature of the data was the presence of both hyperpalmitoylated and depalmitoylated proteins in the STT and LTT trained groups, as well as in their corresponding yoked controls. Clear separation to hyper- and depalmitoylated proteins has previously been reported also after contextual fear conditioning (Nasseri et al., 2022), chronic unpredictable stress (Zareba-Koziol et al., 2019) or induction of synaptic plasticity in several *in vitro* models (Pytyś et al., 2025). In this study, while a small subset of proteins exhibited consistent regulation of palmitoylation across both time points, the majority were uniquely associated with either early or late phases of learning. Interestingly, a fraction of shared proteins displayed opposite directions of palmitoylation, suggesting that dynamic cycles of palmitoylation and depalmitoylation may fine-tune protein function over time. Such bidirectional regulation could reflect transitions between active signaling states and stabilized synaptic configurations, highlighting the reversible nature of this modification as a key regulatory feature. One explanation could be that individual zDHHC palmitoyl acyltransferases and depalmitoylating enzymes act on specific substrates, while their localization and activity are dynamically regulated by synaptic signaling (Abazari et al., 2023; Brigidi et al., 2015; Yokoi et al., 2016). For instance, NMDAR-dependent stimulation drives ZDHHC5 from dendritic spines to shafts, where it palmitoylates δ-catenin on recycling endosomes, promoting its return to spines together with AMPARs (Brigidi et al., 2015). Other zDHHC enzymes (e.g., ZDHHC2, ZDHHC5, and ZDHHC9) are themselves post-translationally modified in an activity-dependent manner, altering their stability and protein interactions (Abazari et al., 2023). ZDHHC2 may be recruited to the postsynaptic density to increase PSD-95 palmitoylation, whereas ZDHHC8 palmitoylates PICK1 during LTD. ZDHHC7 and other family members also display sex- and region-specific synaptic substrate selectivity (Buszka et al., 2023; Woolfrey et al., 2018; Zaręba-Kozioł et al., 2021). Overall, whether proteins become hyper- or depalmitoylated likely depends on the spatial and temporal activation and trafficking of specific zDHHC enzymes and thioesterases, as well as substrate accessibility and crosstalk with other post-translational modifications (e.g., phosphorylation or S-nitrosylation). Future studies are needed to disentangle the intracellular pathways linking external stimuli to selective regulation of palmitoylation in defined groups of proteins.

Surprisingly, several proteins, as well as the direction of their palmitoylation changes, were shared between short-term spatial learning in the present study and contextual fear conditioning reported previously (Nasseri et al., 2022). The partial overlap with proteins identified in contextual fear conditioning paradigms suggests that certain palmitoylation-dependent mechanisms may be conserved across species and learning modalities. In particular, the enrichment of cytoskeletal and synaptic proteins among shared datasets points to a core set of substrates that may underpin early phases of memory formation. For instance, calcium voltage-gated channel auxiliary subunit gamma 8 (TARP γ-8, *Cacng8*) and synaptic vesicle glycoprotein 2A (*Sv2a*) were found to be hyperpalmitoylated following learning [**Figure 3**; (Nasseri et al., 2022)]. TARP γ-8 is preferentially expressed in the hippocampus, where it is important for AMPA receptor abundance and extrasynaptic surface expression (Rouach et al., 2005) whereas CaMKIIα-dependent phosphorylation of TARP γ-8 is required for LTP, learning, and memory (Park et al., 2016). SV2A is a key synaptic vesicle protein, and palmitoylation of synaptic vesicle-associated proteins, including SNAREs, synaptotagmin, and synapsin, is known to regulate exocytosis, endocytosis, and vesicle pool dynamics (B. S. & Talwar, 2025; Matt et al., 2019; Zaręba-Kozioł et al., 2018). However, it remains unclear how palmitoylation of SV2A and TARP γ-8 contributes to learning-related plasticity. Therefore, targeting these proteins in future studies may provide further insight into the molecular mechanisms of learning and memory.

The most striking outcome of our hierarchical analysis was the identification of DAGLA as the only protein whose palmitoylation levels consistently increased with general MWM experience and even more with learning the hidden platform location across the STT and LTT paradigms. DAGLA is the principal neuronal source of the endocannabinoid 2-arachidonoylglycerol (2-AG) and a key regulator of retrograde endocannabinoid signaling at excitatory synapses, where it controls multiple forms of synaptic plasticity through presynaptic CB1 receptor activation (Gao et al., 2010; Katona et al., 2006). Consistent with this role, genetic deletion or pharmacological inhibition of DAGLA impairs hippocampal long-term potentiation and spatial learning in the Morris water maze (Schurman, 2018). The functional consequences of DAGLA palmitoylation remain unknown. We hypothesize that hyperpalmitoylation of DAGLA during spatial learning promotes its membrane association, nanodomain targeting, and localization within postsynaptic signaling complexes while regulating its trafficking and surface turnover (Reisenberg et al., 2012; Zhou et al., 2016). Such changes could enhance the spatial and temporal control of local 2-AG signaling during synaptic plasticity. Whether hyperpalmitoylation alters DAGLA surface retention, endocytic recycling, or enzymatic activity remains to be determined. Nevertheless, our findings identify DAGLA palmitoylation as a previously unrecognized candidate mechanism linking activity-dependent lipid signaling to spatial memory formation, providing a foundation for future mechanistic studies.

This study provides a comprehensive characterization of the rat hippocampal palmitoylome, which has remained largely unexplored. To date, only a few large-scale palmitoylome studies have been performed in rats (Edmonds et al., 2017; Kang et al., 2008; Pytyś et al., 2025), in contrast to the substantially larger body of work available in mice (see Swispalm database). An advancement of this study is the identification of the exact site of modification by palmitate which for a majority of proteins remained unknown. In this study we separately studied palmitoylated proteins or palmitoylated peptides and identified 762 and 1482 proteins, respectively. This difference may arise from the fact that in the protein-level (intact enrichment) workflow, each protein is typically counted once, based on protein-group identification, identification is more conservative (higher confidence thresholds) while proteins with few peptides or low abundance may be missed. In the peptide-level workflow, peptide enrichment reduces suppression by protein size or abundance and even low-abundance proteins can be detected if one modified peptide is captured. Importantly, mapping of cysteine residues substantially expands the current knowledge of the neuronal palmitoylome. We identified exact cysteine positions for palmitate in rat hippocampal proteins and a large proportion of the identified sites have not been previously reported, indicating that S-palmitoylation is far more widespread than currently appreciated. Notably, predominantly, we found 1-2 cysteine residues palmitoylated per protein and only a small fraction of proteins (5.5%) had > 6 cysteine residues palmitoylated. This could suggest that palmitoylation is likely a selective and site-specific regulatory event rather than widespread lipidation across the protein. Attachment of palmitate could thus be a simple on-off mechanism allowing modification to control key properties such as membrane association, subcellular targeting, protein stability, or interactions with signaling partners. This fits the established role of palmitoylation as a dynamic molecular switch. By contrast, only 5.5% of identified palmitoylated proteins contained more than six palmitoylated cysteine sites, indicating that extensive multi-site palmitoylation is relatively uncommon and restricted to specific protein classes. These proteins are often large scaffold, trafficking, or membrane-associated proteins (e.g., clathrin, ankyrin-2, dynein, fatty acid synthase), where multiple palmitoylation sites may provide stronger membrane anchoring, regulate interactions with several compartments, or allow combinatorial control of protein function.

Several limitations of the methods and approaches used in this study should be considered. First, the use of whole hippocampal homogenates precludes cell-type- and subregion-specific resolution, which may mask localized or circuit-specific effects. Second, although ABE-based proteomics enables robust identification of palmitoylated proteins, it does not directly measure the kinetics of palmitoylation turnover. Third, the functional consequences of the observed changes remain to be experimentally validated. Future studies combining cell-type-specific approaches, live imaging, and targeted manipulation of palmitoylation enzymes will be important for establishing causal links between palmitoylation dynamics and synaptic function. Moreover, hydroxylamine-based enrichment captures thioester-linked S-acylation and therefore does not unequivocally distinguish palmitate from other fatty acyl chains (Nederstigt et al., 2025). In addition, site coverage is influenced by peptide detectability, digestion efficiency, protein abundance, and membrane protein recovery; thus, undetected sites cannot be interpreted as unmodified. Localization of the modification may also be less certain for peptides containing multiple cysteine residues. Finally, identification of a modified site does not in itself establish functional relevance and will require targeted biochemical and mutational validation. Despite these limitations, this resource provides a valuable foundation for future mechanistic studies aimed at understanding how site-specific palmitoylation regulates protein function. Given that different cysteine residues within the same protein can have distinct effects on localization and activity, the ability to resolve palmitoylation at the peptide level represents a significant advance.

In summary, our results demonstrate that S-palmitoylation is dynamically and differentially regulated during distinct phases of spatial learning, with early stages characterized by widespread and multifunctional remodeling and later stages by more selective and stabilization-related changes. These findings position S-palmitoylation as a key regulatory mechanism bridging rapid synaptic signaling and long-term structural plasticity. By providing a proteome-wide and site-resolved view of palmitoylation *in vivo,* this study lays the groundwork for future investigations into how this lipid modification shapes neural circuit function and behavior.

## Materials and Methods

### Animals

Two-month-old (250-300 g) male Wistar rats were used for all experiments. Upon arrival at the housing room, the rats were split into cohabiting pairs, and one animal in each pair was marked with a non-toxic marker at the base of the tail to enable individual identification. The animals were housed in an air-conditioned room with monitored temperature (22 ± 1°C) and humidity (50 ± 5%), under a 12/12-h artificial light-dark cycle. The animals had access to drinking water and food *ad libitum*. The rats were handled by the experimenter once a day, for a week before the start of behavioral assays. All experiments were conducted during the dark period of the cycle. The animals were handled according to the Polish guidelines and EU legislation for animal care. All experimental protocols were approved by the local authority for animal experiments (permission number WAW2/019/2024, 2nd Local Ethics Committee for Animal Experiments in Warsaw, Warsaw University of Life Sciences, Warsaw, Poland).

### Morris water maze

The animals were trained in the Morris water maze task (D’Hooge & De Deyn, 2001) to locate a hidden platform in a circular pool with a diameter of 1.5 meters. The pool was assigned four virtual points on its perimeter spaced 90° away from one another labeled as: North (N), East (E), South (S), West (W). The water in the pool was maintained at a temperature of 25-27°C, monitored with a thermometer. A transparent plastic 10 × 10 cm square platform was submerged 1.5 cm below the water surface in the center of the SE quadrant - this quadrant was designated as the target quadrant (TQ). The platform remained in the same position throughout all the spatial learning trials. A dim indirect light was placed over the pool and visual cues (contrasting geometric shapes) were placed on the walls of the room to aid the animals in spatial orientation. The animal’s behavior was recorded with a video camera located above the pool, and using a computerized video-tracking system (Ethovision XT 14, Noldus Information Technologies, Wageningen, The Netherlands). Additional video analysis for spatial search strategy assessment was done in EthoVision XT18. Body-point position data were exported as .csv files and analyzed using the Rtrack software package to determine the spatial search strategy employed by each animal.

### The short-term spatial learning protocol (STT)

A short-term spatial learning training protocol consisted of 15 consecutive trials where the animal learned to locate a hidden platform and a probe trial performed 1 h after the start of the training. Animals were brought into the experimental room 1 h before the start of training to acclimate to the environment. In the training session, the rat was removed from its cage and placed on the platform for 1 min to allow orientation to visual cues. Next, the experimenter introduced the rat into the water, facing the pool wall. In all experiments, the initial starting position of the rat in the maze was at a location opposite to the platform in the OP quadrant. The rat had 1 min to swim and find the platform. If it failed to reach the platform within this time, the experimenter gently guided it to the platform. After spending 1 min on the platform, the rat was placed in the water at one of four starting points (N, E, S, W). The trials alternated between near (S or E) and far (N or W) starting points. The sequence of starting points was kept consistent between the animals. Each rat was subjected to a total of 15 training trials. Each time after the rat was released, the experimenter retreated to an adjacent room where she observed the trial through a live recording from the camera above the pool. The latencies to reach the platform were recorded for each trial. After each trial, the rat spent 1 min on the platform (intertrial interval of 1 min). After completing the training session, the rat was dried and returned to its home cage. For the probe trial, 1 h after the start of the training, the platform was removed from the pool, and the rat was released into the pool at the location opposite to the previous position of the platform. The rat swam for 1 min, after which it was removed from the pool, dried, returned to its home cage, and immediately sent to be sacrificed for tissue collection. The time the rat spent swimming in the target quadrant of the pool and number of crossings into the target zone (corresponding to the prior location of the hidden platform), were recorded as measures of recently acquired spatial memory. The procedure for yoked control animals mimicked the training protocol but omitted the spatial learning aspect of the procedure with the goal of determining which changes in the state of protein S-palmitoylation resulted from swimming activity itself, sensory-motor stimulation, moderate stress, and other factors unrelated to the process of spatial memory acquisition. The visual cues were removed from the walls. In the initial training phase, the yoked control animal was introduced to the pool without a platform 15 times, each time for a duration equal to the average time taken to reach the platform by a paired trained subject (cage-mate), then removed from the pool and placed in a cage outside the pool where it remained for 1 min. The cage location was changed after each trial to prevent the use of olfactory cues for orientation. After 1 h from the start of the swimming session, the rat was placed back into the pool without the platform. The rat swam for 1 min, then was removed from the pool, dried, placed in its home cage, and immediately sent to be sacrificed for tissue collection.

### The long-term spatial learning protocol (LTT)

This protocol was conducted similarly to the described above short-term variant, except that rats underwent four training sessions, one session per day for 4 consecutive days, at the same time each day. Each session consisted of 4 trials, starting once at each of the four starting points (starting point sequence differed between the sessions). The probe trial took place 24 h after the last training session, after which the animals were immediately sent to be sacrificed for tissue collection. Correspondingly, the yoked control animals underwent multi-day swimming sessions analogous to the spatial learning training protocol.

### Acyl-Biotin Exchange Assay

Trained and yoked rats from both the STT and LTT experimental protocols were sacrificed immediately after the probe trial. Cage control animals were sacrificed at a similar time. The acyl-biotin exchange (ABE) assay was used to assess protein S-palmitoylation. Certain anesthetics have been reported to influence lipid raft organization, synaptic protein interactions, and protein S-nitrosylation, which can compete with S-palmitoylation (Agarwal et al., 2022; Krogman et al., 2024; Patel et al., 2020). Therefore, to avoid potential confounding effects of anesthesia on protein palmitoylation, animals were sacrificed by rapid decapitation with no prior anesthesia. Protein extraction from the left rat hippocampi was performed in a Dounce homogenizer in a lysis buffer containing (in mM): 50 Tris-HCl (pH 7.5), 150 NaCl, 1 EDTA, 4% SDS, and 1% Triton X-100. Proteins were reduced using 10 mM TCEP (tris(2-carboxyethyl)phosphine) for 30 min at room temperature (RT). To block free thiol groups resulting from the cysteine reduction, the samples were incubated for 16 h at 4°C with 50 mM N-ethylmaleimide (NEM). To remove unreacted NEM, proteins were then precipitated using the chloroform-methanol method and washed three times with 96% ethanol. The resulting pellets were resuspended in the same lysis buffer. An aliquot of each sample was set aside to produce a pooled-within-condition negative control samples, while the remaining portions of each sample were designated as “positive samples”. To conduct the acyl-biotin exchange reaction, 400 µM of HPDP-biotin (N-[6-(biotinamido)hexyl]-3’-(2’-pyridyldithio)propionamide), a thiol-reactive biotinylation reagent, was introduced to both positive and negative control samples for 1.5 h at RT. However, at the same time, only the positive samples were treated with 1 M hydroxylamine, which served to cleave thioester-linked palmitoyl groups, exposing newly formed thiols to HPDP-biotin. To remove the unreacted compounds, proteins were precipitated using the chloroform-methanol method, washed three times with ethanol, and re-dissolved in an aqueous buffer (in mM: 50 Tris-HCl (pH 7.5), 150 NaCl, 1 EDTA, 0.5% SDS, and 0.2% Triton X-100). For further isolation of the enriched S-palmitoylated protein fraction, equal amounts of protein from each sample were incubated with Pierce High Capacity NeutrAvidin Agarose beads for 2 h at RT. The beads were then washed six times with a buffer (in mM: 50 Tris (pH 7.7), 600 NaCl, and 1 EDTA). To elute the enriched S-palmitoylated proteins, the beads were incubated for 1.5 h at 37°C in 50 mM Tris buffer (pH 7.7) with 1% β-mercaptoethanol. Samples were submitted for Mass Spectrometry analysis to investigate changes in the S-palmitoylation levels of individual proteins.

### Proteomic sample preparation, LC-MS/MS measurements, and data analysis

The collected samples were subjected to chloroform/methanol precipitation and the resulting protein pellets were washed with methanol. The protein was resuspended in 100 mM HEPES pH 8.0 and digested with sequencing grade modified trypsin (Promega) at a protein/enzyme ratio of 100:1 overnight at 37°C [for palmitoylated proteins investigation, in presence of 10 mM TCEP and 15 mM chloracetamide (CAA)]. The digestion was terminated by the addition of a protease inhibitor cocktail (Roche) for palmitoylated cysteines investigation or 1% trifluoroacetic acid (TFA) for palmitoylated proteins investigation. For palmitoylated cysteines investigation, the resulting tryptic peptides were subjected to biotin enrichment using Pierce NeutrAvidin Agarose beads (Thermo Fisher). Briefly, pre-washed beads were incubated with the samples for 2 h RT, washed 3 times with HEPES pH 8.0 and eluted by incubation with 10 mM TCEP, 15 mM CAA in HEPES. The resulting fraction of cysteine containing peptides as well as tryptic peptides for palmitoylated protein investigation were TMT-labeled using on-column protocol <u>(Myers et al., 2019)</u>. Stage tips were packed with three punches of C18 mesh (Affinisep) with a 16-gauge blunt end needle. Resin was conditioned with 150 μl methanol (MeOH), followed by 100 μl 50% acetonitrile (ACN)/0.1% formic acid (FA), and equilibrated with 150 μl 0.1% FA twice. The digest was loaded by spinning at 1200 × g until the entire digest passed through. The bound peptides were washed twice with 150 μl 0.1% FA. 200 μl of TMT reagent in 50 mM phosphate buffer pH 8.0 was loaded over the C18 resin at 300 × g until the entire solution has passed through. The residual TMTs were washed away three times with 150 μl 0.1% FA. Peptides were eluted with 60 μl 60% ACN/0.1% FA. An equal volume of each sample was pooled in one microcentrifuge tube and dried in SpeedVac. To increase the number of proteins identified from complex samples, input (total proteome) samples were additionally fractionated into 8 samples using Pierce High pH Reversed-Phase Peptide Fractionation Kit (Thermo Fisher) according to manufacturers protocol.

Prior to LC–MS/MS analysis, peptides were resuspended in 0.1% TFA and 2% acetonitrile in water. Chromatographic separation was performed on an Easy-Spray Acclaim PepMap column (50 cm × 75 µm i.d., Thermo Fisher Scientific) at 55°C by applying a 120 min acetonitrile gradients in 0.1% aqueous formic acid at a flow rate of 300 nl/min. An UltiMate 3000 nano-LC system was coupled to a Q Exactive HF-X mass spectrometer via an easy-spray source (all Thermo Fisher Scientific). The Q Exactive HF-X was operated in TMT mode with survey scans acquired at a resolution of 60,000 at m/z 200. Up to 15 of the most abundant isotope patterns with charges 2-5 from the survey scan were selected with an isolation window of 0.7 m/z and fragmented by higher-energy collision dissociation (HCD) with normalized collision energies of 32, while the dynamic exclusion was set to 35 s. The maximum ion injection times for the survey scan and the MS/MS scans (acquired with a resolution of 30,000 for palmitoylated cysteines investigation or 45,000 for palmitoylated proteins investigation at m/z 200) were 50 and 120 ms, respectively. The ion target value for MS was set to 3e6 and for MS/MS to 1e5, and the minimum AGC target was set to 1e3.

The data were processed with MaxQuant v. 1.6.17.0 (Tyanova, Temu, & Cox, 2016), and the peptides were identified from the MS/MS spectra searched against Uniprot Rat Reference Proteome (UP000002494) using the built-in Andromeda search engine. For palmitoylated cysteines investigation, cysteine carbamidomethylation was set as a fixed modification and methionine oxidation and alkylation of cysteines with NEM were set as variable modifications. For in silico digests of the reference proteome, cleavages of arginine or lysine followed by any amino acid were allowed (trypsin/P), and up to two missed cleavages were allowed. The FDR was set to 0.01 for peptides, proteins and sites. Unique and razor peptides were used for quantification helping protein grouping (razor peptides are the peptides uniquely assigned to protein groups and not to individual proteins). Other parameters were used as pre-set in the software. Reporter intensity corrected values for protein groups were loaded into Perseus v. 1.6.10 (Tyanova, Temu, Sinitcyn, et al., 2016). Standard filtering steps were applied to clean up the dataset: reverse (matched to decoy database), only identified by site, and potential contaminant protein groups (from a list of commonly occurring contaminants included in MaxQuant) were removed. Reporter intensity corrected values were log_2_ transformed and protein groups with values across all samples were kept. Student’s t-test (1-sided, permutation-based FDR = 0.001, S0 = 1) was performed to identify cysteine-containing peptides with significantly higher intensity values in positive than negative control samples. For palmitoylated protein investigation, raw data of enriched S-palmitoylated protein samples and input (total proteome) samples were processed together using MaxQuant software. Reporter ion MS2-based quantification was applied with reporter mass tolerance = 0.003 Da and min. reporter PIF = 0.75. Cysteine modification by NEM was set as a fixed modification and cysteine carbamidomethylation, methionine oxidation, as well as protein N-terminal acetylation were set as variable modifications. For in silico digests of the reference proteome (trypsin/P), and up to two missed cleavages were allowed. The FDR was set to 0.01 for peptides, proteins and sites. Match between runs was enabled. Other parameters were used as pre-set in the software. Reporter intensity corrected values for protein groups were loaded into Perseus v. 1.6.10. Standard filtering steps were performed as described above. Reporter intensity corrected values were log_2_ transformed and protein groups with min 1 valid value for enriched samples were kept. The values recorded for the enriched samples not treated with hydroxylamine (negative control samples) were subtracted from the values recorded for the enriched samples subjected to hydroxylamine treatment. Values recorded for the input samples were normalized by median subtraction within TMT channels. The values recorded for the input samples were subtracted from the values recorded for the corresponding enriched samples. Median was then subtracted within TMT channels. To determine which proteins were differentially palmitoylated between experimental groups Student’s t-tests (2-sided, permutation-based FDR = 0.05, S0 = 0.1, n = 6 were performed on the groups of PALM samples. In addition, Welch’s t-tests (2-sided, permutation-based FDR = 0.05, S0 = 0.1, n = 6) were performed on the groups of INPUT samples to determine possible difference in protein levels between groups.

The data were exported from Perseus and formatted to its final form in Microsoft Excel 2016.

### Bioinformatic analysis

Functional and protein-protein interaction enrichment analyses, visualization of enrichment networks, and term ranking for DPPs were performed using Metascape (Zhou et al., 2019) (https://metascape.org/gp/index.html#/main/step1). Differentially detected proteins (DPPs) from STT (186 proteins) and LTT (62 proteins) groups were compared against the background input proteome (5,260 proteins). The minimum overlap threshold was set to 3 genes, and significantly enriched terms (p < 0.01, Benjamini–Hochberg false discovery rate corrected) were retained. Only experimentally validated physical interactions were included in network construction. Interactions were retrieved from STRING (physical interaction score > 0.132) and BioGRID. The resulting networks contained proteins with at least one experimentally supported physical interaction within the dataset. Subnetworks comprising 3–500 proteins were further analyzed using the Molecular Complex Detection (MCODE) algorithm to identify densely connected protein clusters, which are presented in the corresponding colored network panels. Further methodological details are available in the Metascape documentation. Additional bioinformatic analyses were performed using custom scripts in R. Protein identifiers were mapped from gene symbols to Entrez Gene IDs using the clusterProfiler package in conjunction with the org.Rn.eg.db. The background universe for enrichment analyses was defined as all proteins identified in the input proteome after removal of missing and duplicate identifiers. KEGG pathway enrichment analyses were performed using clusterProfiler with Benjamini–Hochberg correction, applying an adjusted p-value cutoff of 0.1. For pathway-level visualization enabling integration of quantitative proteomic changes into KEGG pathway topology, log₂ fold-change (log₂FC) values representing palmitoylation differences between trained and yoked animals were mapped to corresponding Entrez Gene IDs for both STT and LTT conditions (https://tomaszwojtowicz-coder.github.io/KEGG/index.html). These data were overlaid onto selected KEGG pathways using the pathview package (organism code: *Rattus norvegicus*, rno), enabling visualization of both magnitude and direction of palmitoylation changes within biologically relevant pathways, including synaptic and neurodegeneration-associated processes. All scripts used for data processing and visualization are publicly available via the RepOD repository (https://doi.org/10.18150/OG5EQV). SwissPalm (Blanc et al., 2015) was used as a reference database for annotations of predicted and experimentally validated palmitoylation sites, as well as for previously reported palmitoylomes.

### Statistical analysis

Behavioral data were analyzed using unpaired Student’s t-tests when comparing two groups (e.g., yoked vs. trained animals). Comparisons of escape latency across repeated behavioral training sessions were analyzed using the Friedman test followed by Wilcoxon signed-rank test with Holm correction for multiple comparisons. Differences in the relative proportions of proteins assigned to subcellular compartments were analyzed using the Chi-square test. Prior to parametric analyses, data distribution normality and homogeneity of variances were assessed using the Shapiro-Wilk test and Brown-Forsythe test, respectively. Data meeting these assumptions were analyzed using parametric tests as indicated above. Principal component analysis (PCA) was performed with Perseus v. 1.6.10 (Tyanova, Temu, Sinitcyn, et al., 2016) for proteomic dataset comparisons to assess sample clustering and variability between experimental groups. All statistical analyses were performed using GraphPad Prism 11 or Python 3.13.9. Statistical significance was defined as *p* < 0.05 and data are presented as mean ± SEM unless otherwise indicated.

## List of supplementary files (https://doi.org/10.18150/OG5EQV)

**Table 1.** Percentages of animals classified into each Rtrack search strategy during successive acquisition trials of the short-term (STT) and long-term (LTT) Morris water maze paradigm (**Supplementary Figure 1**).

**Table 2.** Rat hippocampal homogenate input protein list (5,260 proteins) and results of principal component analysis (PCA) of palmitoylated proteins from naive animals and trained groups compared with their respective time-yoked controls in STT and LTT conditions. Each point represents the palmitoylome profile of an individual animal.

**Table 3.** Palmitoylated proteins detected by ABE-TMT LC-MS/MS (763 proteins, **Figure 2** and **3**, **Supplementary Figure 2**).

**Table 4.** Protein annotation based on UniProt Cellular Component (CC) terms and their distribution (**Figures 2** and **6**).

**Table 5.** Gene Ontology (GO) enrichment analysis of differentially palmitoylated proteins (DPPs). Results of Metascape functional enrichment analysis of hippocampal DPPs following STT and LTT spatial learning protocols.

**Table 6.** KEGG pathways analysis for DPPs identified in STT and LTT groups (**Supplementary Figure 4**).

**Table 7.** Comparison of differentially palmitoylated proteins (DPPs) identified 1 h after contextual fear conditioning in mice (Nasseri et al., 2022) with those identified 1 h after spatial learning (Morris water maze) in rats in the present study **Supplementary Figure 5**).

**Table 8.** List of 1,482 proteins with identified palmitoylated cysteine sites. The table also includes predicted and curated cysteine palmitoylation sites for selected proteins **Figure 6**).

## Supplementary Information

A custom dictionary used to collapse UniProt annotations into 13 major cellular compartments.

## Acknowledgements

This work was supported by the National Science Centre, Poland (grant nr 2019/34/E/NZ4/00387). The authors would like to thank Rafał Czajkowski and Adam Hamed (Nencki Institute of Experimental Biology, Warsaw, Poland) for consultations regarding animal behavior models, and Remigiusz Serwa (Proteomics Core Facility, International Institute of Molecular Mechanisms and Machines of the Polish Academy of Sciences, Warsaw, Poland) for mass spectrometry analysis.

## Author contributions

A.P., R.P., R.K.F., and T.W. conceived and designed the study. A.P., R.I., and T.W. performed the experiments. R.K.F., J.W., and T.W. supervised the study. A.P., R.P., R.K.F., A.C.C.A., L.M.H., A.F.L., R.M., K.R., and T.W. analyzed the data and prepared the figures. K.R. and T.W. wrote the first draft of the manuscript. All authors interpreted the results, critically revised the manuscript, and approved the final version.

